# For common community phylogenetic analyses, go ahead and use synthesis phylogenies

**DOI:** 10.1101/370353

**Authors:** Daijiang Li, Lauren Trotta, Hannah E. Marx, Julie M. Allen, Miao Sun, Douglas E. Soltis, Pamela S. Soltis, Robert P. Guralnick, Benjamin Baiser

## Abstract

Should we build our own phylogenetic trees based on gene sequence data, or can we simply use available synthesis phylogenies? This is a fundamental question that any study involving a phylogenetic framework must face at the beginning of the project. Building a phylogeny from gene sequence data (purpose-built phylogeny) requires more effort, expertise, and cost than subsetting an already available phylogeny (synthesis-based phylogeny). However, we still lack a comparison of how these two approaches to building phylogenetic trees influence common community phylogenetic analyses such as comparing community phylogenetic diversity and estimating trait phylogenetic signal. Here, we generated three purpose-built phylogenies and their corresponding synthesis-based trees (two from Phylomatic and one from the Open Tree of Life [OTL]). We simulated 1,000 communities and 12,000 continuous traits along each purpose-built phylogeny. We then compared the effects of different trees on estimates of phylogenetic diversity (alpha and beta) and phylogenetic signal (Pagel’s λ and Blomberg’s K). Synthesis-based phylogenies generally yielded higher estimates of phylogenetic diversity when compared to purpose-built phylogenies. However, resulting measures of phylogenetic diversity from both types of phylogenies were highly correlated (Spearman’s *ρ* > 0.8 in most cases). Mean pairwise distance (both alpha and beta) is the index that is most robust to the differences in tree construction that we tested. Measures of phylogenetic diversity based on the OTL showed the highest correlation with measures based on the purpose-built phylogenies. Trait phylogenetic signal estimated with synthesis-based phylogenies, especially from the OTL, were also highly correlated with estimates of Blomberg’s K or close to Pagel’s λ from purpose-built phylogenies when traits were simulated under Brownian Motion. For commonly employed community phylogenetic analyses, our results justify taking advantage of recently developed and continuously improving synthesis trees, especially the Open Tree of Life.

## Introduction

Phylogenies describe the evolutionary history of species and provide important tools to study ecological and evolutionary questions (Baum and Smith 2012). Recently, phylogenies have been used to better understand patterns of community assembly. The phylogenetic structure of ecological communities can lend insight into the processes by which local communities assemble from regional species pools (Webb et al. 2002). For example, if closely related species are more likely to co-occur in the same habitats, we might suspect that these species share traits that allow them to have a positive growth rate under the environmental conditions in these habitats. To test whether closely related species are more or less likely to co-occur, one common approach is to calculate the phylogenetic diversity of communities and then compare the observed phylogenetic diversity with those expected by chance through different null models. There is a growing body of literature using this community phylogenetic approach, documenting the phylogenetic structure of ecological communities across taxa and scales (Webb et al. 2002, Cavender-Bares et al. 2006, Helmus et al. 2007, Vamosi et al. 2009, Cardillo 2011, Smith et al. 2014, Li et al. 2017, Marx et al. 2017). Complementing analyses of phylogenetic community structure, phylogenetic signal of ecologically important traits may also be tested (e.g., Cavender-Bares and Reich 2012, Li et al. 2017); traits that have strong phylogenetic signal (i.e., closely related species have more similar trait values than expected by chance) can then provide insights into potential causes of the observed phylogenetic community structure (Webb et al. 2002, Cavender-Bares et al. 2009, Vamosi et al. 2009). Therefore, comparing community phylogenetic diversity and estimating trait phylogenetic signal are two key components of community phylogenetic analyses.

As an important facet of biodiversity, phylogenetic diversity (Faith 1992) also plays a crucial role in conservation biology by complementing more traditional taxonomic measures of biodiversity (e.g., species richness). For example, two communities can have the same number of species but differ drastically in their phylogenetic diversity depending on relatedness of the constituent species. The community with higher phylogenetic diversity, representing taxa more distantly related to each other, is expected to be more stable and productive given its greater evolutionary potential to adapt to changing environmental conditions (Forest et al. 2007, Maherali and Klironomos 2007, Lavergne et al. 2010). Therefore, all else being equal, a community with higher phylogenetic diversity should have higher conservation priority.

The information gained from community phylogenetic analyses is only as good as the species composition data and the phylogenies from which they are generated. In this manuscript, we explore how methods of tree generation affect phylogenetic diversity metrics and phylogenetic signal tests. Generally, ecologists and evolutionary biologists use two common approaches to build phylogenies for community phylogenetic analyses. The first approach is for a researcher to generate his/her own phylogenies for a set of target species based on gene sequence data. We refer to such phylogenies as purpose-built phylogenies. The second approach is to derive phylogenies based on available synthesis trees, such as the Open Tree of Life^1^, or classifications, such as the Angiosperm Phylogeny Group (APG IV et al. 2016), by pruning or sampling, respectively, from the resource so that the phylogeny contains only the target species. We refer to such phylogenies as synthesis-based phylogenies. To a certain extent, one can argue that a synthesis tree could be a purpose-built tree for a larger set of species, but the sources for deriving the synthesis-based trees vary in scope, methodology, assumptions, and content (see Materials and Methods for further description of source trees for synthesis-based phylogenies). From a researcher’s perspective, a purpose-built phylogeny is a major undertaking but offers potential to utilize taxonomic and phylogenetic expertise often needed in order to successfully construct trees. Synthesis trees, as compilations of peer-reviewed phylogenetic hypotheses, offer an immediately available, but typically less customizable output to researchers. We thus use these two terms (purpose-built and synthesis-based) to categorize the underlying methods and researcher cost-benefits to obtain phylogenies.

Generating a purpose-built tree requires more effort, expertise, and cost than subsetting a well-developed phylogeny or sampling from a classification. Generally, purpose-built trees are constructed by using newly generated sequence data and then combining those data with data already available on GenBank, although in many cases the researcher may simply use what is in GenBank. The first step requires gathering tissue for taxa of interest either from field or museum collections, extracting DNA from these tissue samples, and then identifying, amplifying, and sequencing appropriate loci. The gene regions selected are typically based on the taxa of interest and discipline-accepted standards. Resulting sequences are aligned in programs such as MUSCLE (Edgar 2004). Sequences are also commonly sourced entirely or as an addition to sequence data already in databases like GenBank with the help of computational pipelines such as PHLAWD (Smith et al. 2009). Appropriate models of evolution for phylogenetic estimation are determined using programs like PartitionFinder (Lanfear et al. 2012) such that each gene region in a set of concatenated sequences can be treated separately. The most appropriate models of nucleotide evolution are used to estimate phylogenies in Maximum Likelihood (ML) and/or Bayesian Inference (BI) frameworks in programs like RAxML (Stamatakis 2014), MrBayes (Ronquist and Huelsenbeck 2003), and BEAST (Drummond and Rambaut 2007). Depending on the desired application, it may be necessary to impose topological constraints to ease phylogenetic inference or fossil constraints to scale branch lengths to time. Statistics for clade support are calculated using bootstrap or jack-knifing techniques in an ML framework, and posterior probabilities in BI. Despite the fact that multiple software programs are available to help automate these processes (e.g., phyloGenerator (Pearse and Purvis 2013), SUPERSMART (Antonelli et al. 2017)), many decisions at different steps must be made based on expert knowledge (e.g., Which genes to select? How to select models? Which software program to use? How to estimate divergence time?).

Because of the effort, expertise, and cost required to generate purpose-built phylogenies, many community phylogenetic studies use a second approach: deriving phylogenies from available synthesis trees. Over the past few decades, tremendous advances in computational tools and increasingly available genetic sequence data have led to vastly improved phylogenies for plants (Zanne et al. 2014), birds (Jetz et al. 2012), fishes (Rabosky et al. 2013), and mammals (Bininda-Emonds et al. 2007, Fritz et al. 2009). Such advances in phylogenetics have facilitated the synthesis of all available information to make a comprehensive tree of life on Earth (Hinchliff et al. 2015). With these available synthesis trees and software programs such as Phylomatic (Webb and Donoghue 2005), ecologists can derive phylogenies for the species or communities they are interested in with less effort and limited cost. When different studies use the same synthesis tree to derive their phylogenies, their phylogenetic diversity results are comparable. Importantly, this may not be the case if they use purpose-built phylogenies. In addition, these approaches may avoid some issues when generating phylogenies from sequence data such as taxon sampling effects (Park et al. 2018). However, the tractability of phylogenies based on synthesis trees often comes with the cost of decreased resolution (e.g., increase in polytomies) of the resulting phylogenies compared with purpose-built ones; such trees also have taxonomic gaps, which are often filled using existing classifications to become comprehensive.

Previous studies have demonstrated that most phylogenetic diversity (Swenson 2009, Patrick and Stevens 2014, Boyle and Adamowicz 2015) and phylogenetic signal (Molina-Venegas and Rodriguez 2017) metrics are robust to terminal polytomies. These studies, however, used simulated phylogenies or compared different posterior purpose-built phylogenies. Therefore, they provided little practical advice about selecting between purpose-built and synthesis-based phylogenies for ecological studies. In this study, we compared phylogenetic diversity and phylogenetic signal metrics calculated from purpose-built phylogenies and corresponding phylogenies derived from three commonly used sources. It is important to note that we do not treat the purpose-built phylogenies as a gold standard, and we recognize that sampling bias of both taxa and genes, combined with variation introduced through the tree-building process (e.g., tree reconstruction methods, assessment of support, etc.), can compromise the accuracy of purpose-built phylogenies. However, these issues – and others – apply also to the source trees used for synthesis-based phylogenies, although perhaps at different scales. Our aim here is to quantify the influence of the two tree construction techniques on measures of phylogenetic diversity and phylogenetic signal that are commonly employed in the rapidly growing field of community phylogenetics.

## Materials and Methods

### Purpose-built phylogenies

We collected three “purpose-built” phylogenies from published sources. The first purpose-built phylogeny is for 540 plant taxa in the globally critically imperiled pine rockland ecosystem in South Florida, USA (Trotta et al. 2018). The second phylogeny consists of 1,064 alpine plant taxa in France (Marx et al. 2017). The third purpose-built phylogeny has 1,548 plant species with distributions in Florida, USA (Allen et al. 2019). All three phylogenies were estimated from sequence data and were time-calibrated (i.e., chronograms). When using time-calibrated phylogenies, phylogenetic diversity measures the amount of evolution in time-units, and this is the measure we focus on here. For details regarding the phylogenetic tree building processes employed, see the Appendix.

### Commonly available phylogenies

For each of the three purpose-built phylogenies, we generated four phylogenies based on different sources. The first two were generated using Phylomatic v4.2 (Webb and Donoghue 2005) using two different backbone trees: `R20120829` (APG III 2009) and `zanne2014` (Zanne et al. 2014). We call the first phylogeny `tree_apg` and the second one `tree_zanne`. The phylogeny `tree_zanne` has branch lengths because the backbone tree zanne2014 was inferred from seven gene regions for >32k plant species and was time-calibrated using ‘congruification’ (Eastman et al. 2013). In contrast, the phylogeny `tree_apg` has no branch lengths and is based, not on the result of a phylogenetic analysis *per se*, but on a series of phylogenetic analyses as summarized by the Angiosperm Phylogeny Group III (2009). The APG classification is now updated as APG IV (2016), but Phylomatic uses APG III (and the differences between APG III and APG IV are small). To add branch lengths, we used the `bladj` algorithm in Phylocom (Webb et al. 2008) to convert the tree to a chronogram using a set of the minimum node ages given by Wikström et al. (2001).

The third phylogeny was derived from the Open Tree of Life (Hinchliff et al. 2015), a recent comprehensive phylogeny for ~ 2.3 million named species of life, including all eukaryotes, Archaea, and Bacteria. This phylogeny, which we call `tree_otl`, is a supertree constructed from available source trees, with missing species added based on taxonomy; this resulting tree therefore contains many polytomies and does not include branch lengths. To calculate branch lengths, we first identified descendants for each of the internal nodes in `tree_otl` and then searched for their divergence time in the TimeTree of Life database (Kumar et al. 2017). The TimeTree database was compiled based on 3,163 studies and 97,085 species (as of October 10, 2017). For a pair of species included in this database, we extracted their average divergence time from all previous studies. Using the divergence date of internal nodes from the TimeTree database, we then determined branch lengths of `tree_otl` using Phylocom (Webb et al. 2008) and its bladj function. Recently, an updated phylogeny with branch lengths for seed plants based on the Open Tree of Life was published (Smith and Brown 2018); however, we did not use this seed plant phylogeny as a source because it contains only seed plants, and our purpose-built phylogenies also contain other clades of vascular plants.

The fourth phylogeny was a random coalescent phylogeny generated using the rcoal function from the R package `ape` (Paradis et al. 2004). The random tree was then scaled to have a root age that was the average root age of `tree_apg`, `tree_zanne`, and `tree_otl`. Results based on the random phylogeny should not correlate with those based on other phylogenies.

Not every species from the purpose-built phylogenies was found in all of the synthesis phylogenies. For the pine rockland phylogeny, 514 out of 540 species (95.2%) were found in all phylogenies. For the alpine plant phylogeny, 994 out of 1064 species (93.4%) were found in all phylogenies. For the Florida flora phylogeny, 1472 out of 1548 species (95.1%) were found in all phylogenies. Therefore, we pruned the purpose-built phylogenies to have the same species as their corresponding synthesis tree. In practice, one could insert species that were missing from the derived phylogeny as polytomies in the same genus, so that all species could be included in the analysis.

### Generation of community assemblages

For each purpose-built phylogeny, we simulated 1,000 presence/absence site-by-species matrices. Each matrix has 30 sites, with species within each site randomly selected from the phylogeny tips representing the species pool. We fixed species richness of each site to be 50 to remove any effects of species richness on the phylogenetic diversity measures. Without setting all sites to have the same number of species, results based on different phylogenies will correlate with each other. For example, it is likely that results from `tree_random` will be highly correlated with results from other phylogenies (Appendix Fig. A1). This is because most phylogenetic diversity metrics correlate with species richness, which, in turn, will lead to correlations among them and confound the comparisons of effects of phylogeny *per se* on the measurement of phylogenetic diversity. Removing the constraint of using the same species richness does not affect our results and conclusions (Appendix Figs. A1, A2). In our current setting, the maximum total number of species across 30 sites is 30 × 50 = 1500, which is similar to the number of tips in the largest purpose-built phylogeny in our study. We selected species from the species pool randomly because previous studies demonstrated that different approaches to species selection give similar results (Swenson 2009).

### Phylogenetic diversity measurements

For each site-by-species matrix, we calculated alpha and beta phylogenetic diversity for each of the phylogenies using indices that are commonly used in community phylogenetic studies. For phylogenetic alpha diversity, we used Faith’s PD (PD), mean pairwise distance (MPD), and mean pairwise distance between the closest relatives (MNTD). PD calculates the sum of the branch lengths of all species present in an assemblage (Faith 1992). We did *not* include the root of the phylogeny when calculating PD. MPD calculates the average pairwise distance between all species, and MNTD calculates the average pairwise distance between the closest relatives in an assemblage (Webb et al. 2002). We selected these three metrics for phylogenetic alpha diversity among the myriad of metrics available because they are most commonly used and represent different but complementary information about phylogenetic structure of communities (Miller et al. 2017, Tucker et al. 2017).

For phylogenetic beta diversity, we applied UniFrac (Unif), inter-assemblage MPD (MPD_beta), inter-assemblage MNTD (MNTD_beta), and phylogenetic community dissimilarity (PCD) to all possible unique combinations of assemblage pairs. Unif is derived from the Jaccard dissimilarity index and calculates the total branch length unique to each assemblage relative to the total branch length of all species in a pair of assemblages (Lozupone and Knight 2005). Therefore, it measures the fraction of evolutionary history unique to each assemblage. MPD_beta and MNTD_beta were derived from MPD and MNTD, respectively, but instead of comparing species within the same assemblage, they compare species from two different assemblages (Webb et al. 2008). PCD measures pairwise phylogenetic dissimilarity between assemblages by asking how much of the variance of values of a hypothetical trait among species in one assemblage can be predicted by the values of species from another. PCD is independent of species richness of the pair of assemblages and has relatively higher statistical power than other common metrics (Ives and Helmus 2010).

As PD and MNTD are both correlated with species richness (Miller et al. 2017), null models that retain species composition while randomly shuffling tips of the phylogeny are commonly used to standardize phylogenetic diversity results. Despite the fact that MPD is independent of species richness, its variance changes relative to species richness (Miller et al. 2017). Therefore, null models are also frequently applied to MPD. Using the null model, standardized effect size (SES) for each metric can be calculated as 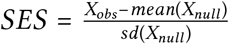, where *X*_*obs*_ is the observed value, and *X*_*null*_ are the *n* values calculated based on null models. Recently, analytic solutions for the SES of phylogenetic alpha diversity metrics were developed (Tsirogiannis and Sandel 2016). The analytic solutions eliminate the need for computationally expensive simulations used to calculate SES values, especially for studies in high-diversity systems. In our simulations, because all sites have the same species richness, we expected that the SES values based on the analytic solutions would have the identical results as the observed phylogenetic diversity values for the statistical analyses we conducted (correlation and linear mixed models, see the Statistical analyses section below). Our simulations confirmed this expectation (Appendix Fig. A3-A6). No analytic solutions for the SES of Unif, MNTD_beta, and PCD are available. However, the pairwise beta diversity metrics share the same core formula with their corresponding alpha diversity metrics. We thus expect that the results based on SES of these beta diversity metrics will be the same as those based on the observed diversity values in our simulations. Given the similarity in results between raw and standardized phylogenetic alpha diversity measures and the large computational burden of calculating SES for phylogenetic beta diversity metrics, we did not include the results for SES in this study.

### Traits simulation and phylogenetic signal

For each purpose-built phylogeny, we simulated continuous traits with two common models of evolution: Brownian Motion (BM) and Ornstein-Uhlenbeck (OU). For both evolution models, we set the rate of trait divergence (sigma, *σ*^2^, a scaling term) to one of three values: 0.2, 0.75, and 1.5. For the OU model, we further varied the strength of selection (alpha, *α*) to be one of three values: 0.05, 0.5, and 1. Note that if alpha = 0, the OU model becomes the BM model. We simulated 12 (3 *σ*^2^ × 4 *α* levels) continuous traits for each purpose-built phylogeny. For each simulated trait, we then estimated its phylogenetic signal with all 5 phylogenies using Pagel’s lambda (λ) (Pagel 1999) and Blomberg’s K (Blomberg et al. 2003), two methods that are most widely used in ecology. Both λ and K have expected values of 1 if a trait evolved along the phylogeny under a BM evolution model. We repeated this process 1,000 times, resulting in 180,000 estimates of phylogenetic signal (3 datasets × 3 sigma × 4 alpha × 5 phylogenies × 1,000 replicates). For traits that were simulated under the BM model (i.e., alpha = 0), we expected that the average values of both estimated λ and K to be 1 when tested with the purpose-built phylogenies. For traits that were simulated under strong OU models (alpha = 0.5 and 1 here), we expected the average values of both estimated λ and K to approach zero (i.e., weak signal), regardless of which phylogeny we used. Note that K can approach, but will never be, zero by definition. In addition, we examined the type I error rates (i.e., false positive) in estimating λ and K for all phylogenies by randomly reshuffling trait values that were simulated under the BM model with *σ*^2^ = 0.2, resulting in another 15,000 estimates of phylogenetic signal (3 datasets × 5 phylogenies × 1,000 replicates).

### Statistical analyses

We have three primary goals. First, we want to test the correlation between phylogenetic diversity values calculated from purpose-built phylogenies and those calculated from synthesis-based phylogenies. For this goal, we calculated the average Spearman’s rank-based measure of the correlation between phylogenetic diversity values from all phylogenies across the 1,000 simulations. We used rank-based correlation because we are interested in relative, rather than absolute, phylogenetic diversity.

Second, we want to investigate whether phylogenetic diversity calculated from synthesis-based phylogenies over- or under-estimates phylogenetic diversity when compared to purpose-built phylogenies. For this goal, we used Linear Mixed Models (LMMs) with phylogenetic diversity values from the purpose-built phylogeny as the response variable, the phylogenetic diversity values from one of the synthesis-based phylogenies as the predictor, and the simulation dataset as the random term. We scaled the diversity values to have mean zero and standard deviation one before fitting the models. We also forced the regression line through the origin. If the slope of the regression line is significantly different from zero, then phylogenetic diversity based on purpose-built phylogenies and synthesis-based phylogenies is significantly correlated. Furthermore, if the slope is higher/lower than one, then the phylogenetic diversity values based on the synthesis-based phylogenies are lower/higher than those based on the purpose-built phylogeny. For pairwise beta diversity, because one site can be compared with all other sites, the beta diversity values are not independent. To account for this, we included datasets, `site1` within each dataset (the first site in the site pair), and `site2` within each site (the other site in the site pair) as random terms in the LMMs (cf. Li and Waller 2017).

Third, we want to determine which synthesis-based phylogeny estimated phylogenetic signal values that are the closest to those estimated with the purpose-built phylogeny. For this question, we mostly relied on data visualization instead of statistical tests because of the large sample size (n = 1,000). Furthermore, Pagel’s λ had very small variances when estimating with the purpose-built phylogenies (< 10^−7^ for all simulations under BM); such small variances led all estimated correlation coefficients to be around zero. Thus, we only focus on the absolute differences in the estimated λ values between the purpose-built phylogeny and the synthesis-based phylogenies. For Blomberg’s K, we compared estimated values of `tree_purpose` with those from other synthesis-based phylogenies using Spearman’s rank correlations. We used non-parametric tests for Blomberg’s K because it has a highly skewed distribution. The workflow of this study is outlined in Fig. 1. All analyses were conducted with R v3.4.3 (R Core Team 2017).

**Figure 1:**
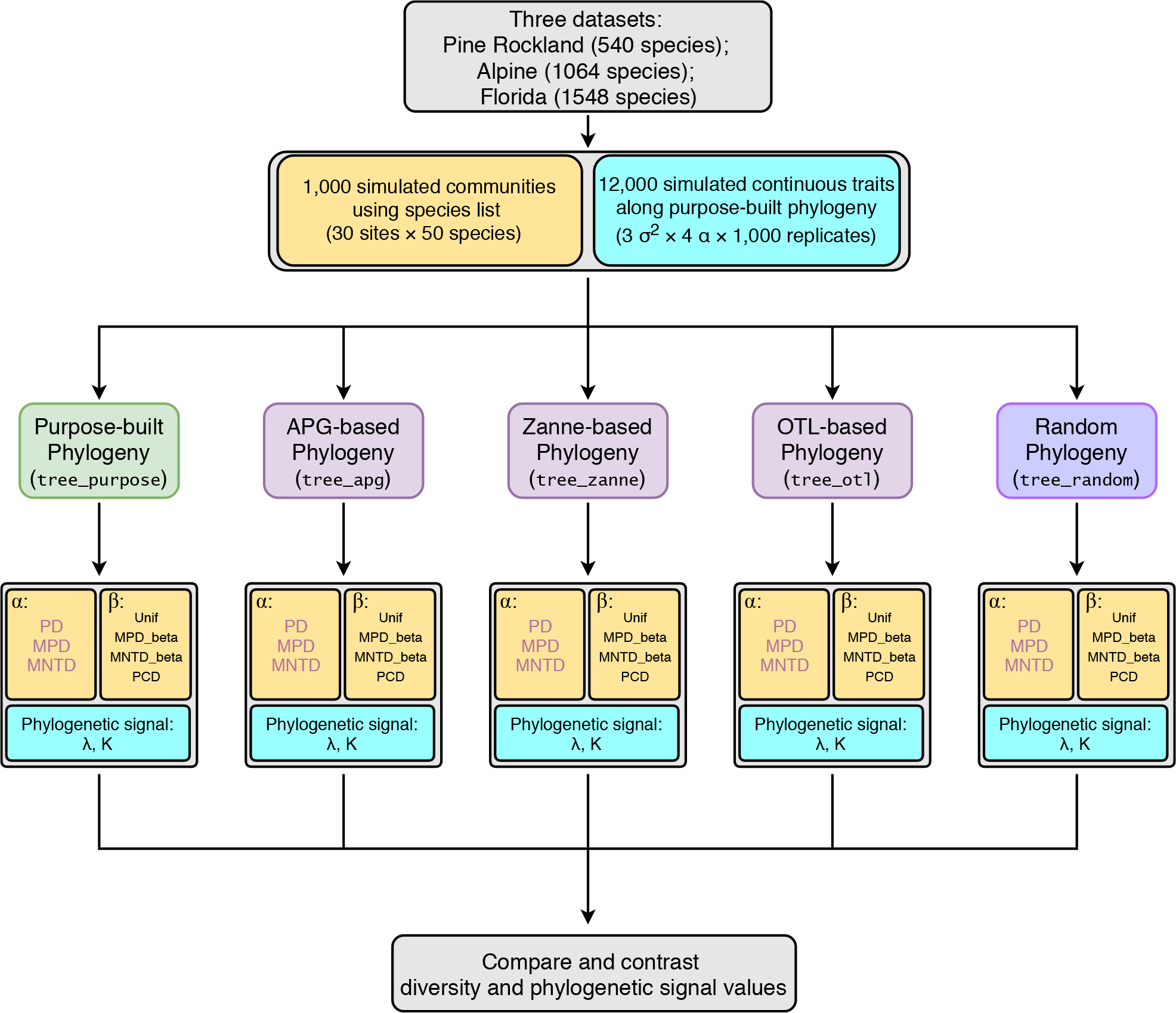
Workflow to assess effects of commonly used synthesis phylogenies on community phylogenetic diversity and trait phylogenetic signal estimations. Boxes with light yellow background are related to community phylogenetic diversity; boxes with light blue background are related to trait phylogenetic signal. Abbreviations: APG, Angiosperm Phylogeny Group; OTL, Open Tree of Life; PD, Faith’s Phylogenetic diversity; MPD, Mean pairwise distance; MNTD, Mean nearest taxon distance; Unif, Unifraction; PCD, Phylogenetic community dissimilarity; λ, Pagel’s lambda; K, Blomberg’s K.

## Results

### Alpha diversity

Phylogenetic alpha diversity (PD, MPD, and MNTD) values calculated with different phylogenies (`tree_purpose`, `tree_apg`, `tree_zanne`, and `tree_otl`) were highly correlated. The median Spearman’s correlation of the 1,000 simulations was larger than 0.63 across all comparisons (p < 0.05 for all simulations and comparisons; Fig. 2). In most cases, the median Spearman’s correlation was larger than 0.85, especially for PD and MPD. Therefore, PD and MPD were more robust to varying the source of the phylogeny than MNTD. Across all comparisons, diversity values based Compare and contrast diversity and phylogenetic signal values on `tree_otl` showed the highest correlations with those based on `tree_purpose`, with an average correlation across all comparisons of 0.902. As expected, diversity values based on the random phylogeny `tree_random` were not correlated with diversity values based on other phylogenies, with median Spearman’s correlations close to zero (Fig. 2).

**Figure 2:**
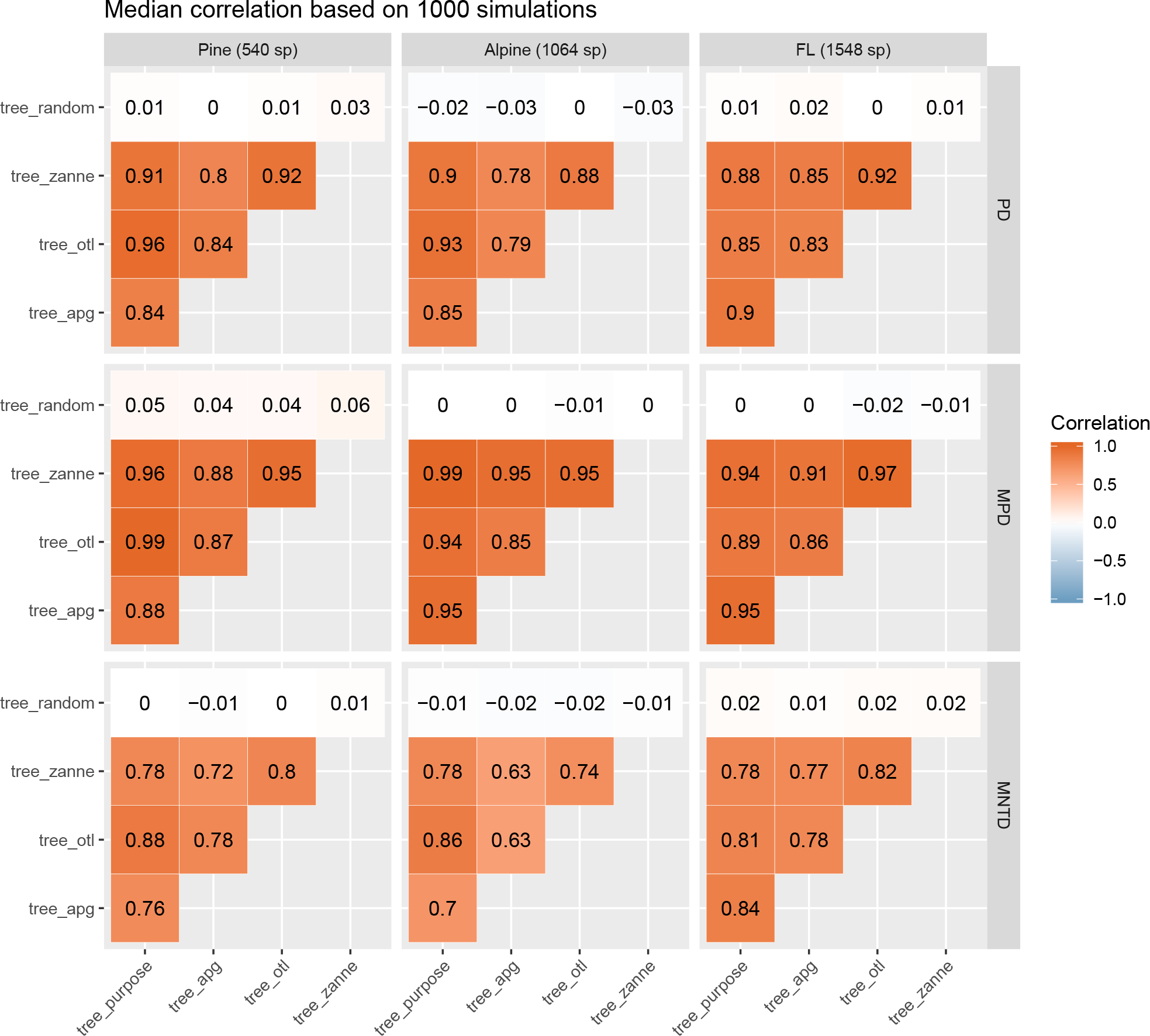
Median correlations of phylogenetic alpha diversity values based on different phylogenies.

The slopes of linear mixed models (LMM) were all less than one (Table 1), suggesting that diversity values based on synthesis-based phylogenies generally were higher than the diversity values based on the purpose-built phylogenies. The PD metrics based on the Open Tree of Life phylogeny (`tree_otl`) had estimates closest to those calculated from the purpose-built phylogenies (Table 1).

**Table 1:** Slopes based on linear mixed models (LMMs). Within the model, the response variable is the phylogenetic alpha diversity values based on the purpose-built phylogeny; the predictor is the phylogenetic alpha diversity values based on one of the synthesis-based phylogenies (`tree_apg`, `tree_zanne`, `tree_otl`, and `tree_random`). Therefore, slopes less than one indicate that diversity values based on synthesis-based phylogenies were higher than those based on the purpose-built phylogenies. Numbers within parentheses are the 95% confidence intervals for the slopes.

| index | dataset | tree_apg | tree_zanne | tree_otl | tree_random |
| --- | --- | --- | --- | --- | --- |
| PD | Pine (540 sp) | 0.843 (0.837, 0.849) | 0.917 (0.913, 0.922) | 0.971 (0.969, 0.974) | -0.001 (-0.013, 0.01) |
| PD | Alpine (1064 sp) | 0.854 (0.848, 0.86) | 0.915 (0.91, 0.919) | 0.937 (0.933, 0.941) | -0.022 (-0.034, -0.01) |
| PD | FL (1548 sp) | 0.92 (0.916, 0.924) | 0.891 (0.886, 0.896) | 0.871 (0.865, 0.876) | 0.006 (-0.005, 0.018) |
| MPD | Pine (540 sp) | 0.891 (0.885, 0.896) | 0.972 (0.969, 0.974) | 0.996 (0.995, 0.997) | 0.047 (0.036, 0.059) |
| MPD | Alpine (1064 sp) | 0.957 (0.954, 0.96) | 0.997 (0.997, 0.998) | 0.941 (0.937, 0.945) | 0.004 (-0.008, 0.015) |
| MPD | FL (1548 sp) | 0.962 (0.958, 0.965) | 0.95 (0.946, 0.953) | 0.895 (0.889, 0.9) | -0.002 (-0.014, 0.009) |
| MNTD | Pine (540 sp) | 0.78 (0.773, 0.788) | 0.787 (0.78, 0.794) | 0.897 (0.892, 0.902) | 0.006 (-0.006, 0.017) |
| MNTD | Alpine (1064 sp) | 0.713 (0.705, 0.721) | 0.794 (0.787, 0.801) | 0.874 (0.869, 0.88) | -0.016 (-0.028, -0.004) |
| MNTD | FL (1548 sp) | 0.856 (0.85, 0.862) | 0.797 (0.79, 0.804) | 0.831 (0.824, 0.837) | 0.03 (0.018, 0.041) |

### Beta diversity

The phylogenetic beta diversity results (Unfi, MPD_beta, MNTD_beta, and PCD) show a similar pattern to the alpha diversity results. Beta diversity of community pairs based on different phylogenies was also highly correlated, with the median Spearman’s correlation from the 1,000 simulations greater than 0.69 across all comparisons (Fig. 3). Overall, phylogenetic beta diversity is more sensitive to the source of the phylogeny than alpha diversity. MPD_beta is the most robust beta diversity metric to the source of the phylogeny, followed by MNTD_beta, Unif, and PCD. Again, PD metrics based on `tree_otl` showed the highest correlation with metrics based on the purpose-built tree, followed by `tree_zanne` and `tree_apg`. Beta diversity values based on `tree_random` did not correlate with values based on any other phylogeny.

**Figure 3:**
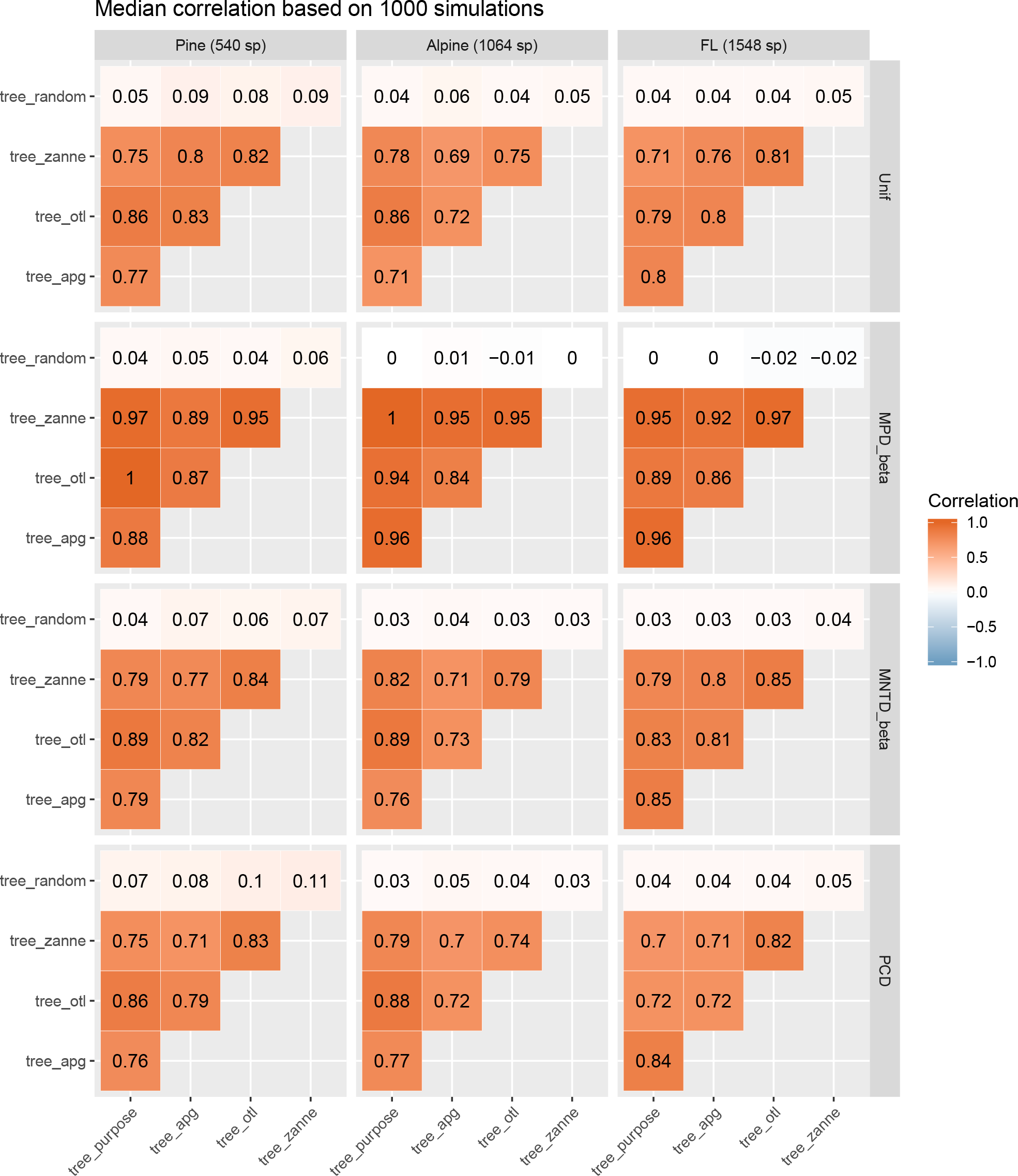
Median correlations of phylogenetic beta diversity values based on different phylogenies.

The slopes of LMMs were generally less than one (Table 2), suggesting that beta diversity values based on synthesis-based phylogenies also were higher than the diversity values based on the purpose-built phylogenies. However, slopes for MPD_beta values based on `tree_otl` were all greater than one, suggesting that beta PD metrics were lower than those calculated from the purpose-built trees. Metrics based on `tree_zanne` for the flora of Florida dataset were also lower than those calculated from the purpose-built tree (Table 2). For the other beta diversity metrics (i.e., Unif, MNTD_beta, and PCD), `tree_otl` generally gave results closer to those based on the purpose-built trees than did the other synthesis-based phylogenies.

**Table 2:** Slopes based on linear mixed models (LMMs). Within the model, the response variable is the phylogenetic beta diversity values based on the purpose-built phylogeny; the predictor is the phylogenetic beta diversity values based on one of the synthesis phylogenies (`tree_apg`, `tree_zanne`, `tree_otl`, and `tree_random`). Therefore, slopes less than one indicate that diversity values based on synthesis-based phylogenies were higher than those based on the purpose-built phylogenies. Numbers within parentheses are the 95% confidence intervals for the slopes.

| index | dataset | tree_apg | tree_zanne | tree_otl | tree_random |
| --- | --- | --- | --- | --- | --- |
| Unif | Pine (540 sp) | 0.824 (0.822, 0.826) | 0.791 (0.789, 0.793) | 0.87 (0.869, 0.872) | 0.063 (0.058, 0.067) |
| Unif | Alpine (1064 sp) | 0.811 (0.808, 0.813) | 0.871 (0.869, 0.873) | 0.896 (0.894, 0.897) | 0.056 (0.053, 0.06) |
| Unif | FL (1548 sp) | 0.871 (0.869, 0.873) | 0.791 (0.788, 0.793) | 0.814 (0.812, 0.816) | 0.071 (0.066, 0.075) |
| MPD_beta | Pine (540 sp) | 0.34 (0.337, 0.342) | 0.972 (0.969, 0.975) | 1.23 (1.225, 1.234) | 0.009 (0.007, 0.011) |
| MPD_beta | Alpine (1064 sp) | 0.797 (0.794, 0.799) | 0.976 (0.976, 0.977) | 1.122 (1.117, 1.127) | 0.002 (0.001, 0.004) |
| MPD_beta | FL (1548 sp) | 0.778 (0.776, 0.781) | 1.343 (1.339, 1.347) | 1.805 (1.797, 1.813) | 0.001 (-0.001, 0.002) |
| MNTD_beta | Pine (540 sp) | 0.856 (0.853, 0.859) | 0.857 (0.854, 0.86) | 0.928 (0.926, 0.93) | 0.054 (0.05, 0.058) |
| MNTD_beta | Alpine (1064 sp) | 0.896 (0.894, 0.899) | 0.952 (0.95, 0.954) | 0.942 (0.94, 0.943) | 0.046 (0.043, 0.05) |
| MNTD_beta | FL (1548 sp) | 0.787 (0.785, 0.789) | 0.762 (0.76, 0.764) | 0.75 (0.748, 0.752) | 0.039 (0.036, 0.043) |
| PCD | Pine (540 sp) | 0.857 (0.854, 0.86) | 0.828 (0.825, 0.831) | 0.872 (0.87, 0.875) | 0.089 (0.085, 0.093) |
| PCD | Alpine (1064 sp) | 0.827 (0.825, 0.83) | 0.912 (0.909, 0.915) | 0.907 (0.905, 0.909) | 0.059 (0.055, 0.063) |
| PCD | FL (1548 sp) | 0.802 (0.799, 0.804) | 0.744 (0.741, 0.746) | 0.719 (0.716, 0.722) | 0.054 (0.05, 0.059) |

### Phylogenetic signal

For all simulated traits, estimated phylogenetic signal (both Pagel’s λ and Blomberg’s K) of `tree_random` were all around 0 as expected (Fig. A7). Therefore, we excluded those values from the comparisons. The divergence rate (*σ*^2^) did not affect the results (Figs. A8, A9). Therefore, we only focus here on *σ*^2^ = 0.2.

Estimated Pagel’s λ values of `tree_otl` were the closest to those of `tree_purpose` among all three synthesis-based phylogenies for both the pine rockland and alpine datasets (Fig. 4) when traits were simulated under BM and weak OU (alpha = 0.05). For the Florida dataset, this is not the case when traits were simulated under BM. Here, average estimated Pagel’s λ values of `tree_apg` were slightly closer to the expected value than `tree_otl`. However, `tree_apg` had much larger variance (Fig. 4) and lower log likelihood (Fig. A10) compared with `tree_otl`. Therefore, `tree_otl` had the best fit among all three synthesis-based phylogenies. The absolute differences of average estimated Pagel’s λ values between `tree_purpose` and `tree_otl` were small when traits were simulated under BM (< 0.022 in all datasets) or weak OU (< 0.13 in all datasets). Furthermore, estimated Pagel’s λ values of `tree_otl` were all significantly different from 0 when traits were simulated under BM and weak OU (high statistical power, Table A1). Together, these results suggest that `tree_otl` can provide relatively close estimates of Pagel’s λ values, has high statistical power, and controls type I error well (Table A1).

**Figure 4:**
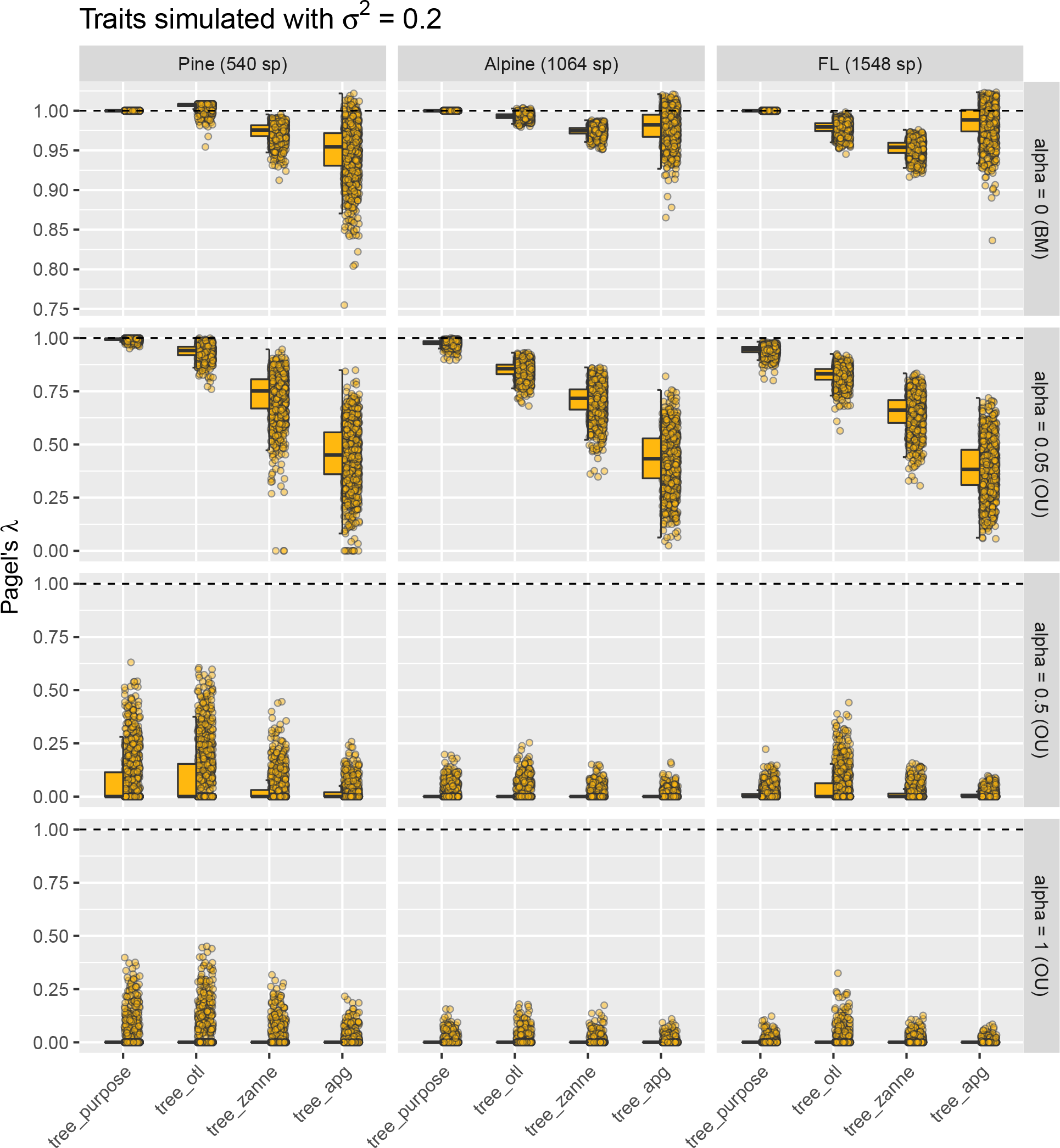
Estimated Pagel’s λ for traits simulated with divergence rate *σ*^2^ of 0.2. When traits were simulated under BM and weak OU models, estimated Pagel’s λ values based on `tree_otl` were the closest to those estimated based on `tree_purpose` in most cases and had smaller variances than other synthesis-based phylogenies. Note that we allow λ to be larger than 1 in all estimates.

For traits simulated under BM, the average values (not the median by definition) of estimated Blomberg’s K of `tree_purpose` were all about 1 as expected (Fig. 5). However, the estimated values had large variance (standard deviation > 0.7) and were skewed (Fig. 5). The high variance allowed us to compare estimated K values between `tree_purpose` and the three synthesis-based phylogenies statistically. When traits were simulated under BM, estimated K values of synthesis-based phylogenies were all significantly different from those estimated with `tree_purpose` (except `tree_apg` for the alpine dataset, paired Wilcoxon tests). However, their values were highly correlated with those estimated with `tree_purpose` (all Spearman’s *ρ* > 0.9, p ≪ 0.001, Fig. 6). When traits were simulated under weak OU (alpha = 0.05), estimated K values of `tree_otl` have the highest Spearman’s *ρ* (all > 0.7) with those of `tree_purpose` and the highest statistical power compared to other synthesis-based phylogenies (Table A1). Compared to Pagel’s λ, Blomberg’s K has higher statistical power when traits were simulated under OU (Table A1). All phylogenies had good type I error controls when estimating phylogenetic signal with Blomberg’s K (Table A1). Together, these results suggest that `tree_apg` can provide relatively close estimates of Blomberg’s K values when the number of species is small. When the number of species is large (e.g., > 1,500), both `tree_otl` and `tree_apg` work well.

**Figure 5:**
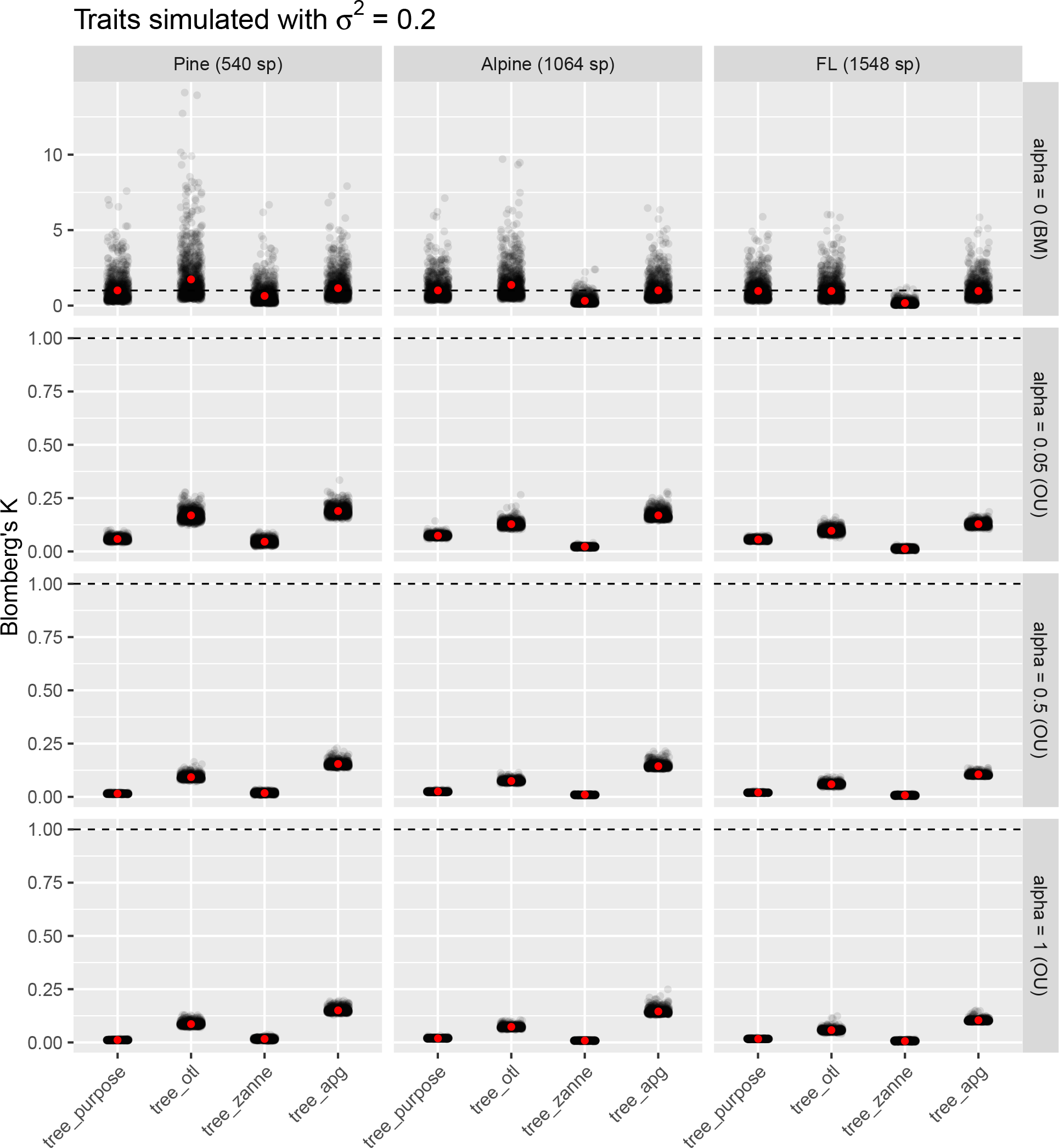
Estimated Blomberg’s K for traits simulated with divergence rate *σ*^2^ of 0.2. Because for Blomberg’s K, it is the mean, not the median, value that has the expected value of 1, we did not use boxplots as in Fig. 4. Instead, we added the average values (red points) on top of jittered raw estimated values.

**Figure 6:**
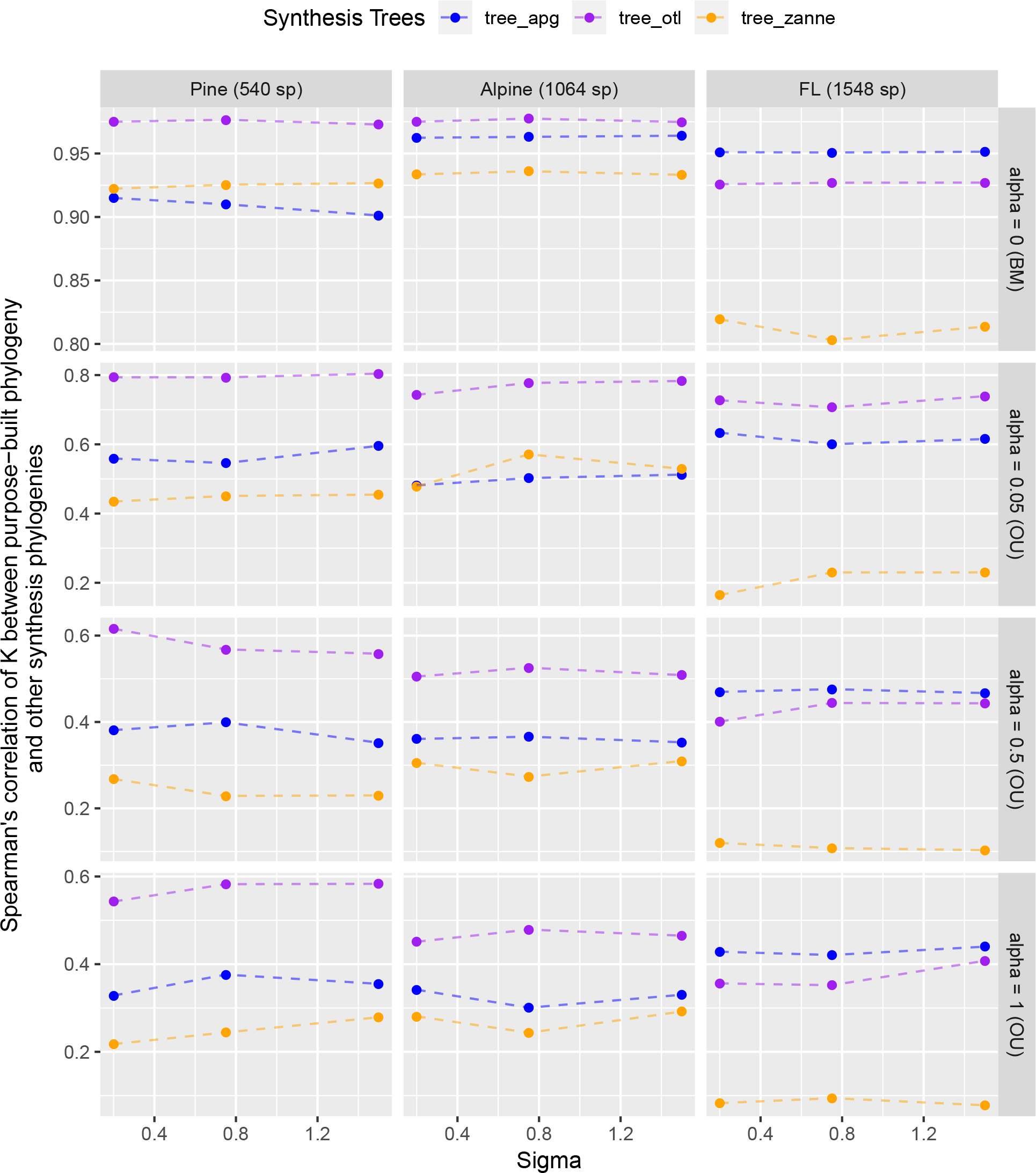
Spearman’s rank correlations of estimated Blomberg’s K values between `tree_purpose` and the three synthesis-based phylogenies.

## Discussion

We examined how different phylogenies, purpose-built and synthesis-based, influenced phylogenetic diversity measures (alpha and beta) and trait phylogenetic signal commonly used in community phylogenetic analyses. We found three main results. First, the synthesis-based phylogenies generally yield higher estimates of phylogenetic diversity compared with purpose-built phylogenies. This is not surprising because synthesis-based phylogenies generally have higher proportions of polytomies than purpose-built ones, which, in turn, leads to larger distances between species within these polytomies. This result agrees with Boyle and Adamowicz (2015) and Qian and Zhang (2016) but contradicts Swenson (2009), who found that phylogenies with more polytomies under-estimated phylogenetic diversity. Second, phylogenetic diversity values calculated from synthesis trees were highly correlated with those based on purpose-built phylogenies, even if the former were higher. These results hold for both alpha and beta diversity and for phylogenies with different numbers of tips. Third, estimated Pagel’s λ values of `tree_otl` were very close to expected values when traits were simulated under BM or weak OU. Estimated Blomberg’s K values of `tree_otl` had high correlation (Spearman’s *ρ* > 0.9) with expected values when traits were simulated under BM. While our study focuses on plants, we expect that our results will generalize to any taxonomic group. Therefore, phylogenies derived from synthesis trees, especially from the Open Tree of Life, can provide similar results to purpose-built phylogenies while saving effort, time, and cost when quantifying and comparing phylogenetic diversity of communities and the phylogenetic signal of traits.

As ecologists and conservation biologists, we mostly care about the relative diversity among communities instead of their absolute diversity. For example, for a set of communities within one region, we may be interested in which communities have the highest/lowest phylogenetic diversity. The absolute phylogenetic diversity of each community does not mean much without comparing it to other communities. Because phylogenetic diversity values based on different phylogenies are highly correlated with each other, the information available for community phylogenetic questions does not differ much between approaches. Even though such synthesis-based phylogenies may yield higher absolute phylogenetic diversity for communities, the relative phylogenetic diversity among communities will be similar to those calculated from typically better resolved but more difficult to obtain purpose-built phylogenies. Based on the information provided by relative values of phylogenetic diversity, the potential improved resolution of purpose-built trees for calculating the absolute PD may not be worth the effort for community phylogenetic questions.

Our finding that phylogenetic diversity metrics are relatively insensitive to the phylogenies from which they are derived has been supported by other recent studies. For example, using simulated fully bifurcating and gradually unresolved phylogenies, Swenson (2009) found that phylogenetic diversity measures are generally robust to the uncertainty of the phylogenies, especially if the uncertainty is concentrated in recent nodes of the phylogeny. Using multiple posterior phylogenies of bats, Patrick and Stevens (2014) rearranged branches across these phylogenies and also found that phylogenetic diversity measures are robust to the phylogenies from which they are calculated. More recently, Cadotte (2015) transformed a phylogeny with different evolution models and found that phylogenetic diversity measures are insensitive to the branch lengths of the phylogeny; getting the topology right is more important when calculating phylogenetic diversity. Qian and Zhang (2016) found similar phylogenetic diversity values of the angiosperm tree flora of North America based on phylogenies derived from Zanne et al. (2014) and Phylomatic (Webb and Donoghue 2005). These studies, however, only focused on alpha diversity. Our study extends the literature by also examining the effects of phylogenies on beta diversity. We found the same pattern for beta diversity and alpha diversity. Taken together, a general pattern emerges: community phylogenetic alpha and beta diversity metrics are robust to reasonably good modern phylogenies.

Why are phylogenetic diversity values from purpose-built and synthesis-based phylogenies highly correlated? There are two possible reasons. First, both purpose-built and synthesis phylogenies likely share a similar systematic backbone and empirical resources such as genes, taxonomies, and expert knowledge. This guarantees that phylogenetic diversity based on these phylogenies will not be dramatically different. Second, phylogenetic diversity metrics aggregate (by summing or averaging) all information into one value for each site, which could help buffer most uncertainty and further mask most of the differences between different phylogenies.

Our results for trait phylogenetic signal suggest that synthesis-based phylogenies can be used as reasonable proxies for purpose-built phylogenies in estimating phylogenetic signal. In our simulations, synthesis-based phylogenies can either slightly overestimate (`tree_otl`), underestimate (`tree_zanne`), or produce largely unbiased estimates (`tree_apg`) of trait phylogenetic signal when the phylogeny is small (< 1,000 species). However, estimated values based on synthesis-based phylogenies were either highly correlated with (Blomberg’s K) or close to (Pagel’s λ) those estimated from the “true” phylogeny (`tree_purpose`) under the BM trait evolution model. A recent study that suggested Pagel’s λ is more robust to polytomies and suboptimal branch-length information in the phylogeny than Blomberg’s K (Molina-Venegas and Rodriguez 2017). Furthermore, another previous study found that Blomberg’s K overestimated phylogenetic signal if a phylogeny has a large proportion of polytomies (Davies et al. 2012). Traits in these studies, however, were simulated only under the BM model of evolution. Our simulations of traits under the OU model of evolution suggested that, compared to Pagel’s λ, Blomberg’s K is more sensitive (more changes in estimated values when alpha changed from 0 to 0.05) and has higher statistical power in identifying less-than-BM phylogenetic signal, making it a more sensitive tool to detect departures from the BM model (Münkemüller et al. 2012). This might be because Blomberg’s K is more sensitive to the pattern of covariances generated by the OU model of evolution than is Pagel’s λ. Therefore, our results suggest that both Pagel’s λ and Blomberg’s K should be used in identifying phylogenetic signal given their own strength and weakness.

Our results should encourage ecologists to increasingly include phylogenetic analyses in community ecology studies, given the growing accessibility of synthesis-based phylogenies and the robustness of phylogenetic diversity and phylogenetic signal measures based on them. Compared with purpose-built phylogenies, synthesis-based phylogenies generally have broader taxon sampling coverage, use more fossil calibration points, and reflect up-to-date taxon classifications. Therefore, we expect synthesis-based phylogenies to be more accurate in terms of topology and node ages, which some have argued are more important than branch lengths for phylogenetic diversity estimation (Cadotte 2015). However, our results should not discourage the construction of purpose-built phylogenies, which are clearly valuable for many ecological and evolutionary questions. This is especially the case for purpose-built trees constructed from local DNA samples. The sequencing of species in a given community can yield data for species that have never been sequenced before. These new sequences can then be incorporated into synthesis trees, improving their resolution for future research. Direct sequencing of samples collected for a community is also important when the community contains undescribed (Pons et al. 2006) or cryptic species (Hebert et al. 2004). Furthermore, for many taxonomic groups, synthesis trees are not available or are far too poorly sampled, and constructing purpose-built trees is the only approach possible for community phylogenetic analyses.

## Conclusion

Community phylogenetics is rapidly becoming an important component of community ecology, macroecology, and biodiversity conservation (Webb et al. 2002, Vamosi et al. 2009). For calculations and comparisons of community phylogenetic diversity and trait phylogenetic signal, an important question arises: can we derive phylogenies from already-available synthesis trees, or should we generate our own purpose-built phylogenies? Our results suggest that phylogenies derived from common synthesis trees yield higher estimates of phylogenetic diversity metrics when compared to purpose-built trees, but values of phylogenetic diversity are highly correlated with those of purpose-built trees. Furthermore, estimated trait phylogenetic signal using synthesis-based phylogenies was reasonably close to (Pagel’s λ) and had high correlations with (Blomberg’s K) expected values based on the purpose-built phylogenies. Particularly, the Open Tree of Life, which includes all major phylogenetic groups (e.g. plants, birds, fishes, mammals, insects, fungi, Archaea, Bacteria), produced the most similar values of community phylogenetic diversity and trait phylogenetic signal when compared to metrics derived from purpose-built trees. Furthermore, a recently updated Open Tree of Life phylogeny for seed plants has branch lengths calculated based on molecular data (Smith and Brown 2018). With new data and studies continuously being integrated into synthesis trees such as the Open Tree of Life, these resources are poised to continue to improve rapidly. As a result, for common community phylogenetic analyses such as comparing phylogenetic diversity among communities and estimating trait phylogenetic signal, we recommend taking advantage of recent well-developed products such as the Open Tree of Life.

## Data Accessibility

All phylogenies and R code used will be uploaded to figshare upon acceptance.

## Acknowledgments

We thank Anthony R. Ives, three anonymous reviewers, and editor Jeannine Cavender-Bares for constructive comments that have greatly improved this manuscript. This study was supported by NSF grants ABI-458034 to BB, DEB-1442280 and DBI-1458640 to PSS and DES, EF-1115210 and DBI-1547229 to PSS, and EF-1550838 (supported HEM).

## Footnotes

1 https://tree.opentreeoflife.org/opentree

