## Supplemental information for "For common community phylogenetic analyses, go ahead and use synthesis phylogenies"

### Appendix S1

*Daijiang Li, Lauren Trotta, Hannah E. Marx, Julie M. Allen, Miao Sun, Douglas E. Soltis, Pamela S. Soltis, Robert P. Guralnick, and Benjamin Baiser*

*15 April, 2019*

**Manuscript title**: For common community phylogenetic analyses, go ahead and use synthesis phylogenies

**Journal**: *Ecology*

### Section S1: details about the three purpose-built phylogenies used

#### Pine Rockland community phylogeny

The pine rockland community phylogeny was constructed from a combination of field and herbarium based collections and supplemented with pre-existing sequences from the Flora of Florida project at the University of Florida. The Floristic Inventory of South Florida (FISF), which includes a comprehensive list of pine rockland species (n = 592, Gann GD 2018), guided our sampling efforts. Specimens were collected in fruit or flower over the course of all seasons and across multiple pine rockalnd fragments in Miami Dade County from 2014 to 2016. We collected material from 331 new field collections, 17 herbarium collections at Fairchild Tropical Botanic Garden (FTBG; Miami, FL, USA) or the University of Florida (FLAS; Gainesville, FL, USA), and 58 field-collected plants currently in cultivation at Fairchild Tropical Botanic Garden. For field and herbarium collections, we extracted total genomic DNA and amplified three commonly used plastid barcodes (Kress et al. 2009): *rbcL*, *matK*, and *psbA-trnH*. Amplification success was confirmed with gel electrophoresis and Sanger sequencing was performed by Beckman Coulter Genomics (Cambridge, Massachusetts, USA), Genewiz (South Plainfield, New Jersey, USA), or Eurofins Genomics (Louisville, Kentucky, USA). In all, we were able to include 540 taxa in our community phylogeny, 90.88% of all vascular plants that occur on the FISF species list.

Both newly generated and pre-existing sequences were edited and assembled using Geneious R9 (Biomatters, Auckland, New Zealand), and resulting alignments were constructed using the MAFFT v1.3.5 (Katoh and Standley 2013) plugin in Geneious. We visually inspected individual alignments before concatenating all three loci into a final alignment. We used PartitionFinder v1.1.1 (Lanfear et al. 2012) and the Akaike information criterion (AIC) to determine the optimal model of evolution for each barcode as well as the best overall partitioning scheme, which were used for all down stream Maximum likelihood (ML) and Bayesian Inference (BI) analyses. We conducted all analyses on the HiPerGator 2.0 supercomputing cluster at the University of Florida. ML analysis was performed in RAxML v8.2.8 (Stamatakis 2014) with an ordinal level topological constraint, following the Angiosperm Phylogeny Group IV (APG IV et al. 2016) for flowering plants and the Pteridophyte Phylogeny Group (Schneider et al. 2016) for ferns and lycophytes. One thousand bootstrap replicates were used to determined clade support. We used a single lycophyte species (*Selaginella eatonii*) as the outgroup. Dating analysis was performed in BEAST 2.4.6 (Bouckaert et al. 2014). Our BEAST analysis included nine fossil constraints selected from previous molecular dating studies in addition to ordinal and superclade constraints sensu APG IV (Schuettpelz and Pryer 2009, Bell et al. 2010, Magallón et al. 2013, APG IV et al. 2016). The tree topology that resulted from our ML search was used as the starting tree for this analysis after it was calibrated to match the fossil constraints using the `chronos` function in the R package ape (Paradis et al. 2004). More details about all included analyses can be found in Trotta et al. (2018).

#### Alpine community phylogeny

For each of the species occurring in the Écrins National Park, France (n = 1,345), we used the PHLAWD pipeline (Smith et al. 2009) to retrieve sequence data for five gene regions (*atpB*, *rbcL*, *matK*, *trnTLF*, and *ITS*) from GenBank (release 209). Sites that were missing across >50% of the taxa in the output alignment were removed from each gene region using PHYUTILITY (Smith and Dunn 2008), and maximum likelihood (ML) inference implemented in RAxML (Stamatakis 2014) was used to estimate gene trees. Outlier taxa (i.e. those falling outside of clades defined by the APG III taxonomy) were identified by visual inspection of each gene tree and were removed before concatenation into a final alignment using PHYUTILITY (Smith and Dunn 2008). The resulting super-matrix consisted of 79% (n = 1,065) of the species surveyed and was used to estimate a ML community phylogeny of the Écrins flora in RAxML v8.0.4, under a GTR-CAT model of nucleotide evolution with simultaneous rapid bootstrap and ML search using 999 replicates, which is ideal for large nucleotide alignments (Stamatakis 2014). We used the 'congruification' approach (Eastman et al. 2013) to map divergence times from a reference time-tree (Soltis et al. 2011, Zanne et al. 2014) with concordant nodes on the best ML estimate of tree topology in the R package `geiger` v2.0.6 (Pennell et al. 2014), and the phylogeny was scaled to time using treePL (Smith and O'Meara 2012). Further details can be found in Marx et al. (2017).

#### Florida Flora phylogeny

We constructed a phylogeny for the vascular flora of Florida based on sequence data of two plastid genes commonly used in plant phylogenetics: *rbcL* and *matK*. We first leveraged existing published sequence data on GenBank. We obtained sequence data from GenBank for 463 species. We checked these sequences throughout the alignment and tree-building process and removed problematic sequences. The remaining species lacked published DNA sequence data. We thus collected fresh materials of these species and sequenced them and deposited voucher specimens at the University of Florida herbarium (FLAS). For rare and hard-to-access species, we sampled specimens from FLAS. We aligned sequences of *rbcL* and *matK* according to a reference protein-coding sequence using Pal2Nal (Suyama et al. 2006). We used PartitionFinder v1.1.1 and AIC (Lanfear et al. 2012) to determine the optimal model of evolution. We then ran RAxML v7.03 (Stamatakis 2014) on the partitioned dataset. We used 100 bootstrap replicates to determine the internal support for the initial tree evaluation. We used three outgroup taxa using data from GenBank: *Physcomitrella patens* (Funariaceae), *Syntrichia ruralis* (Pottiaceae), and *Mastigophora woodsii* (Mastigophoraceae). We examined all phylogenetic trees for issues including possible misidentifications of species, contaminants, and misplaced taxa. We used the best tree from RAxML as a starting tree in a Bayesian analysis using MrBayes v3.2.2 (Ronquist et al. 2012) for $10^6$ generations with 20 independent runs. The top 10 trees were sampled based on likelihood score as representations of the posterior distribution of trees. We used the tree with the highest likelihood in this study. We dated the tree, producing a chronogram, using r8s (Sanderson 2003) and 17 calibration points with a maximum age of 377 million years. More details about the phylogeny building process can be found in Allen et al. (2019).

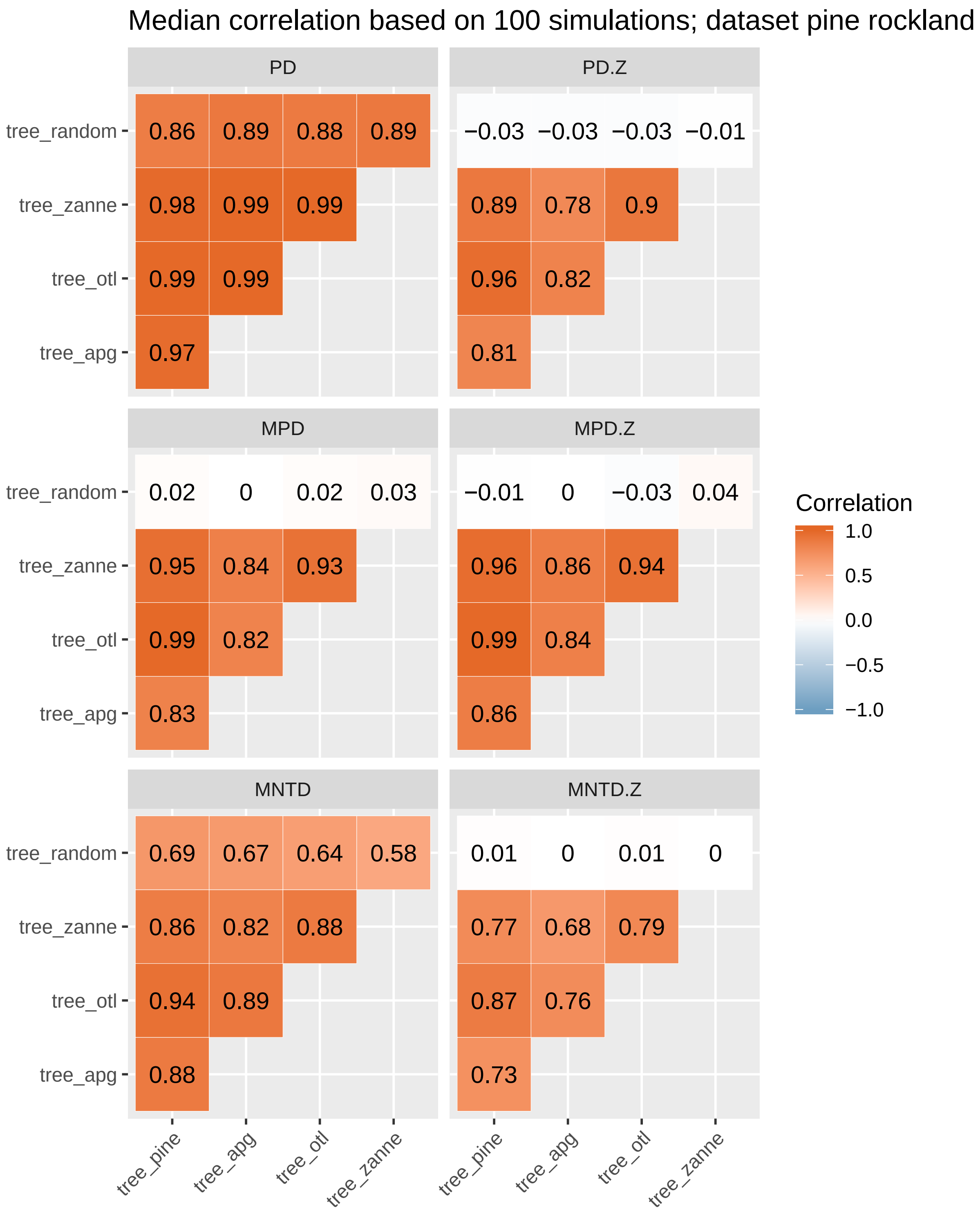


Figure **S1**: Median correlations of phylogenetic alpha diversity values based on 100 simulations of the pine rockland dataset. Within each simulation, different sites have *different* species richness. Results are similar to those in the main text, where species richness was the same across all sites within each simulation to remove effects of species richness. Note that `PD` and `MNTD` (but not `MPD`) are correlated with species richness, thus the high correlations between `tree_random` and other phylogenies. Null models successfully removed these correlations (`PD.Z` and `MNTD.Z`).

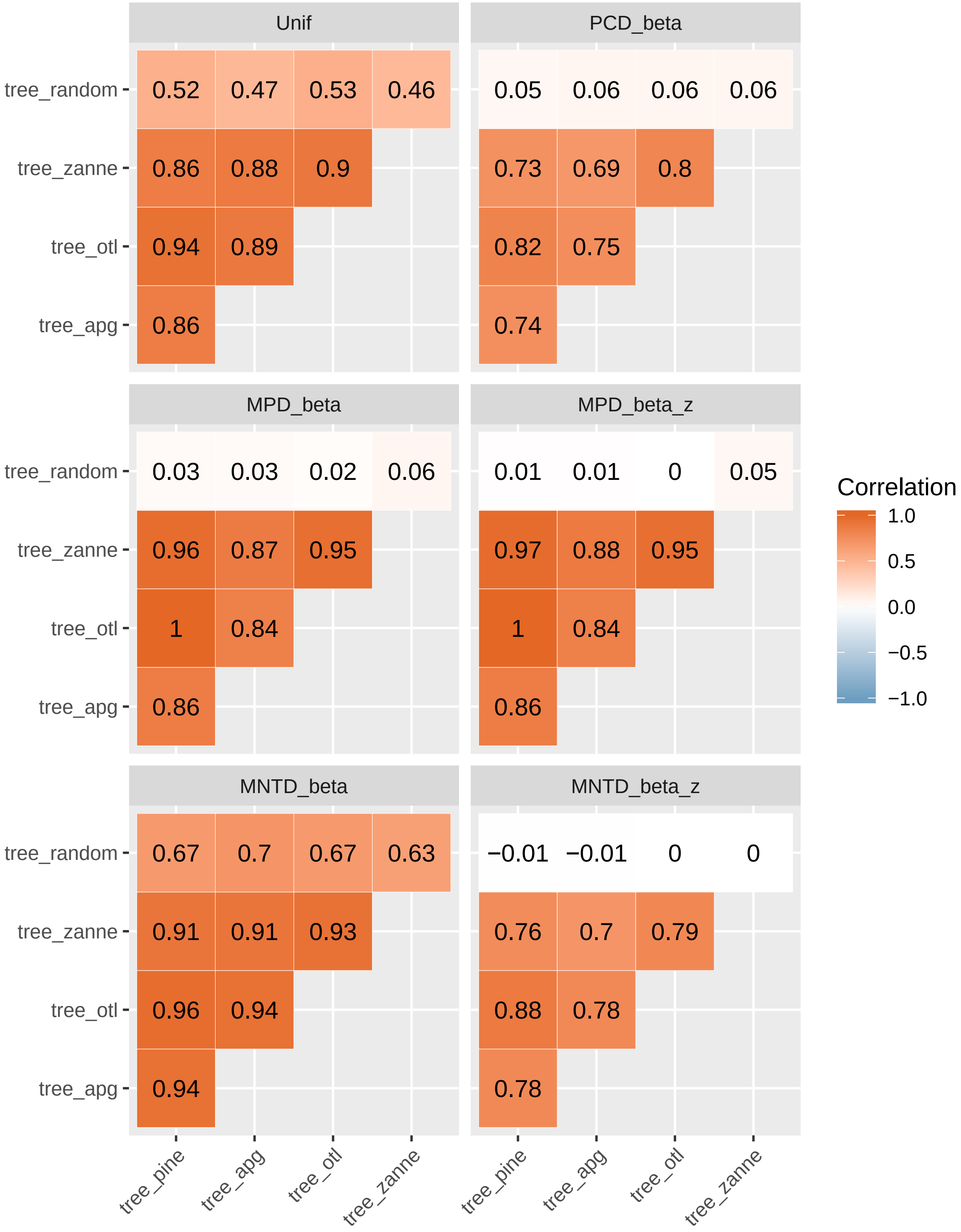


FIGURE **S2**: Median correlations of phylogenetic beta diversity values based on 100 simulations of the pine rockland dataset. Within each simulation, different sites have *different* species richness. Results are similar to those in the main text, where species richness was the same across all sites within each simulation to remove effects of species richness. Note that `Unif` and `MNTD_beta` (but not `MPD_beta` and `PCD_beta`) are correlated with species beta diversity, thus the high correlations between `tree_random` and other phylogenies. Null models successfully removed these correlations (`MPD_beta_z` and `MNTD_beta_z`).

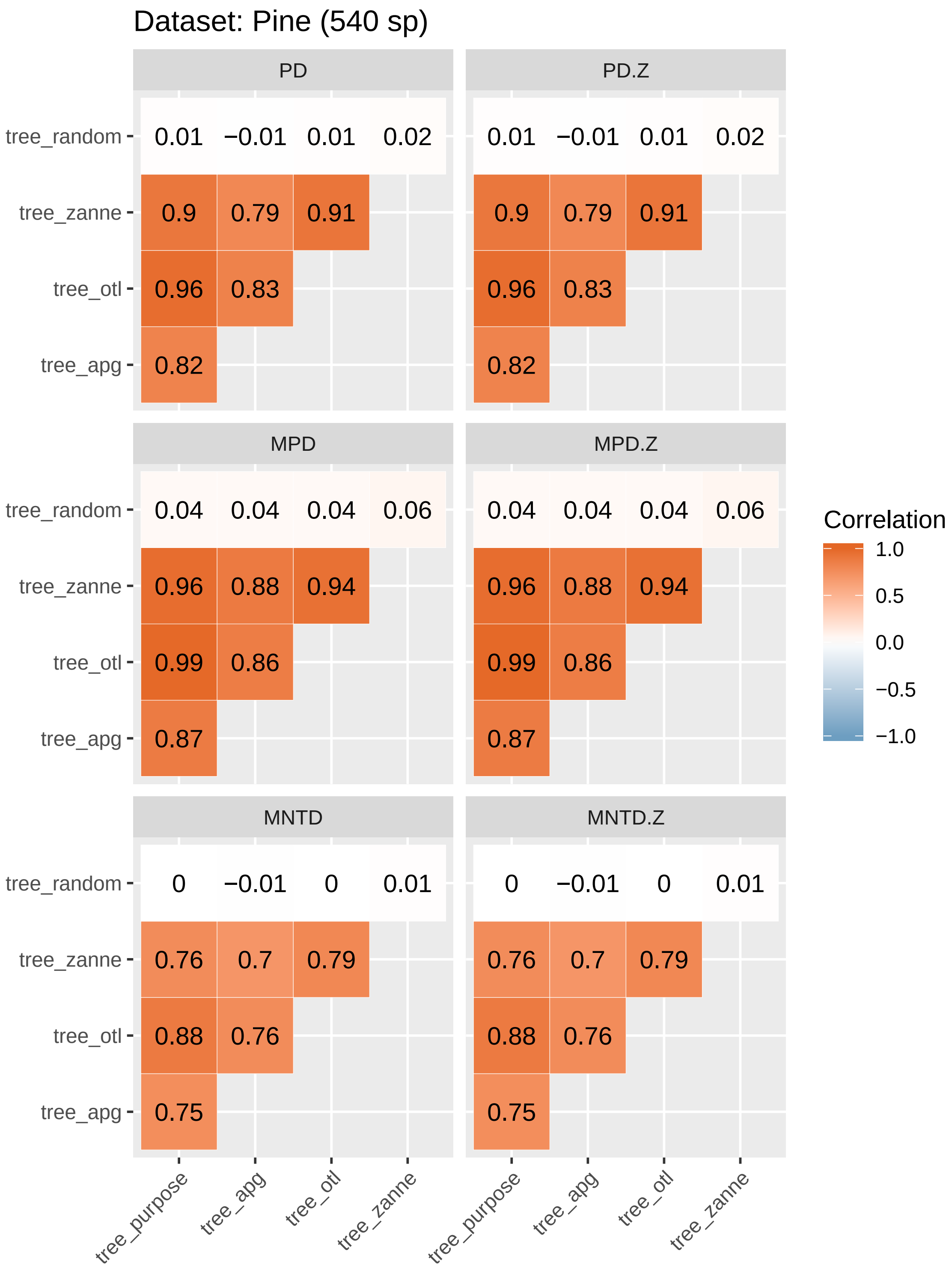


FIGURE S3: Correlations among observed phylogenetic alpha diversity values calculated from different phylogenies for the pine rockland dataset are the same as correlations among their corresponding standardized effect size (SES) calculated from different phylogenies.

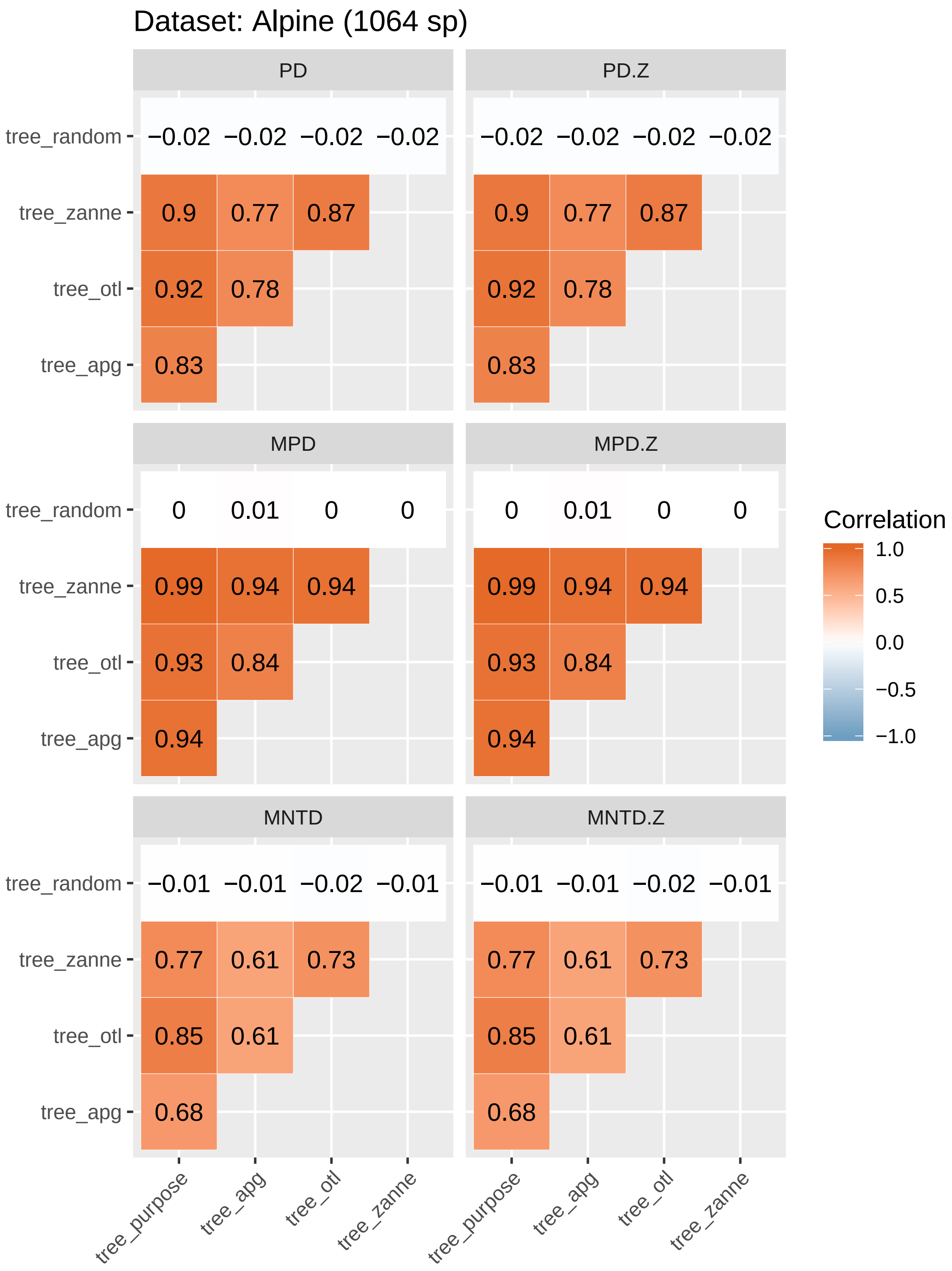


Figure **S4**: Correlations among observed phylogenetic alpha diversity values calculated from different phylogenies for the alpine dataset are the same as correlations among their corresponding standardized effect size (SES) calculated from different phylogenies.

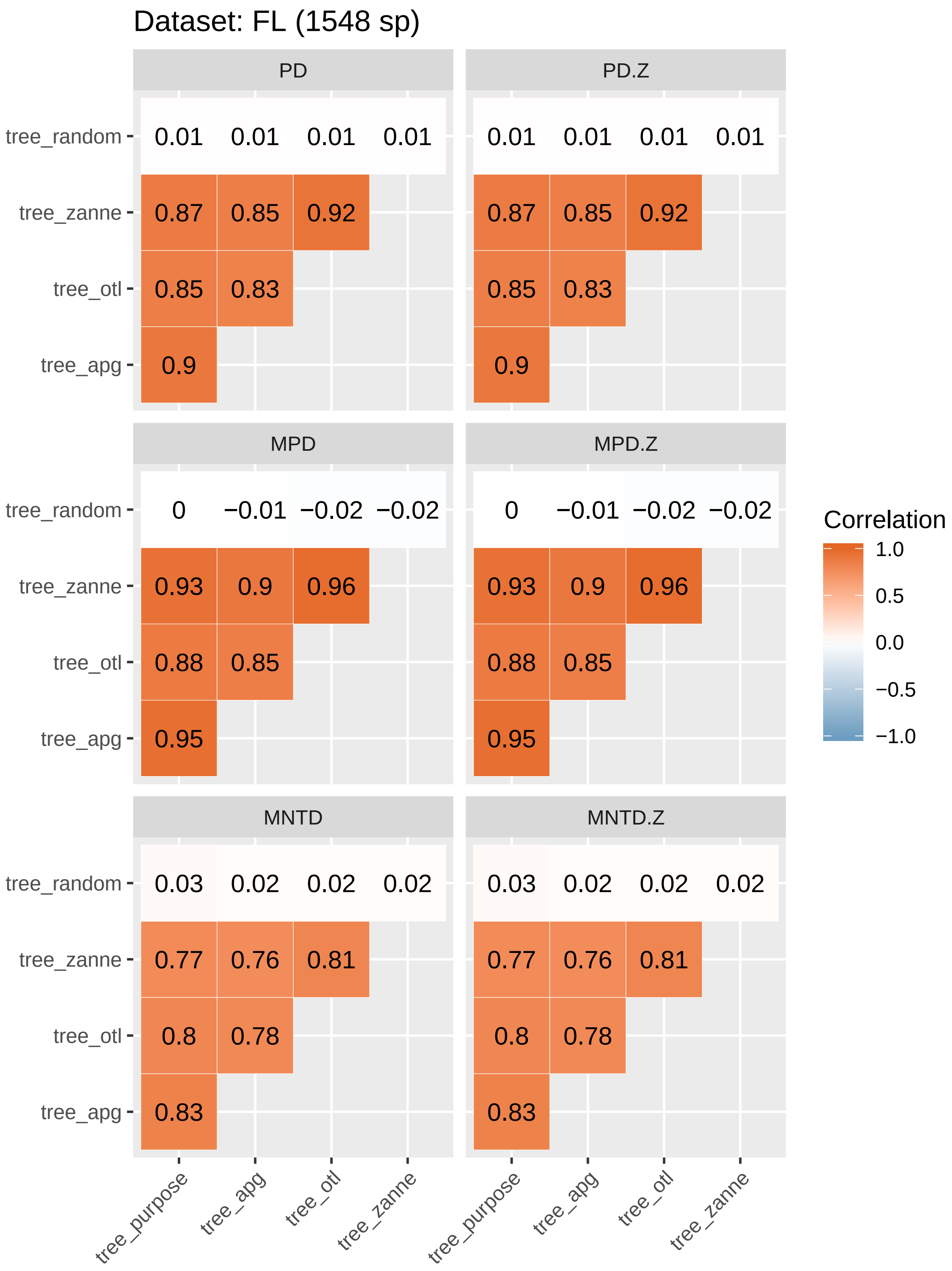


Figure **S5**: Correlations among observed phylogenetic alpha diversity values calculated from different phylogenies for the Florida flora dataset are the same as correlations among their corresponding standardized effect size (SES) calculated from different phylogenies.

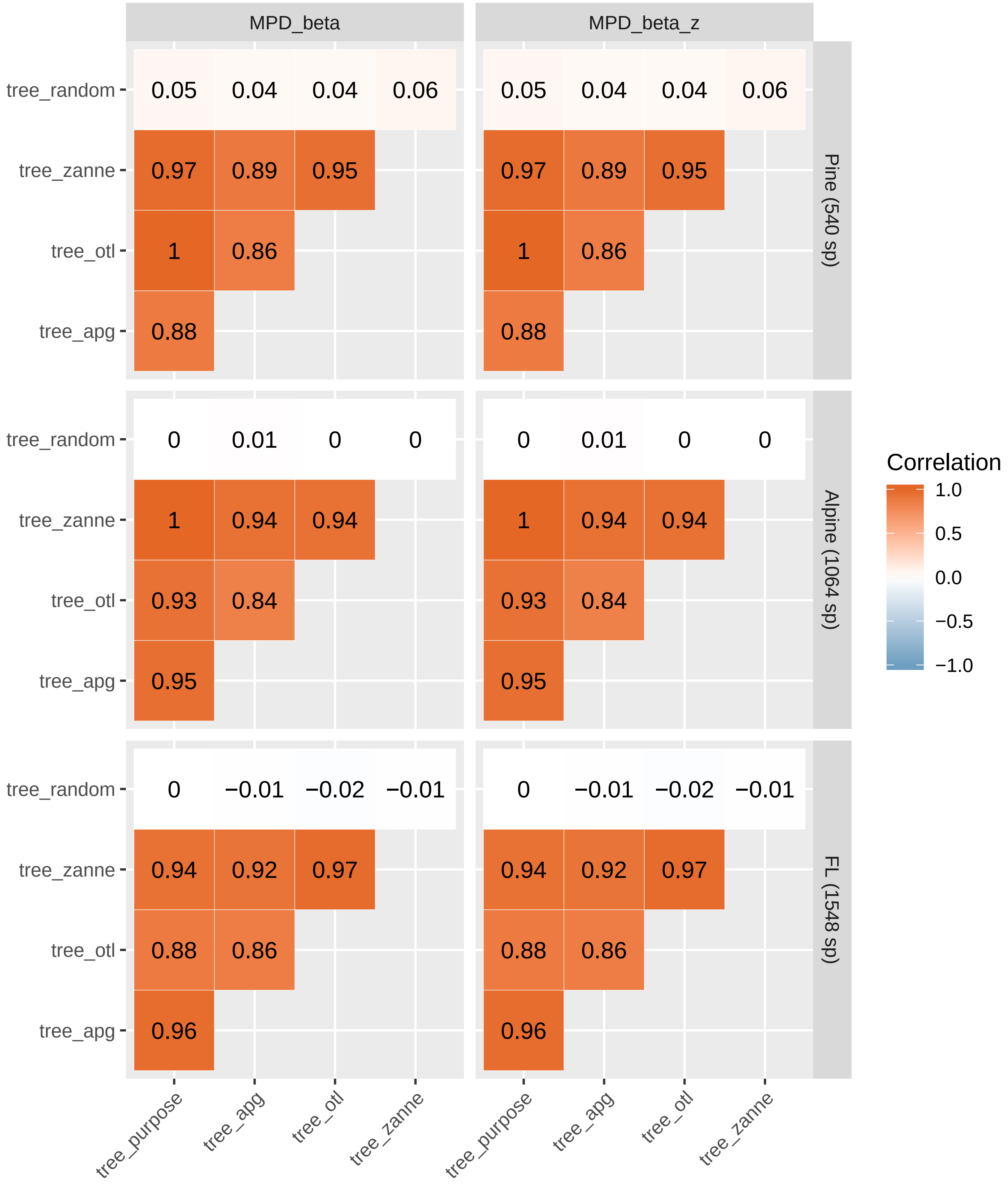


Figure S6: Correlations among observed phylogenetic beta diversity (`MPD_beta`) values calculated from different phylogenies for all datasets are the same as correlations among their corresponding standardized effect size (SES, `MPD_beta_z`) calculated from different phylogenies.

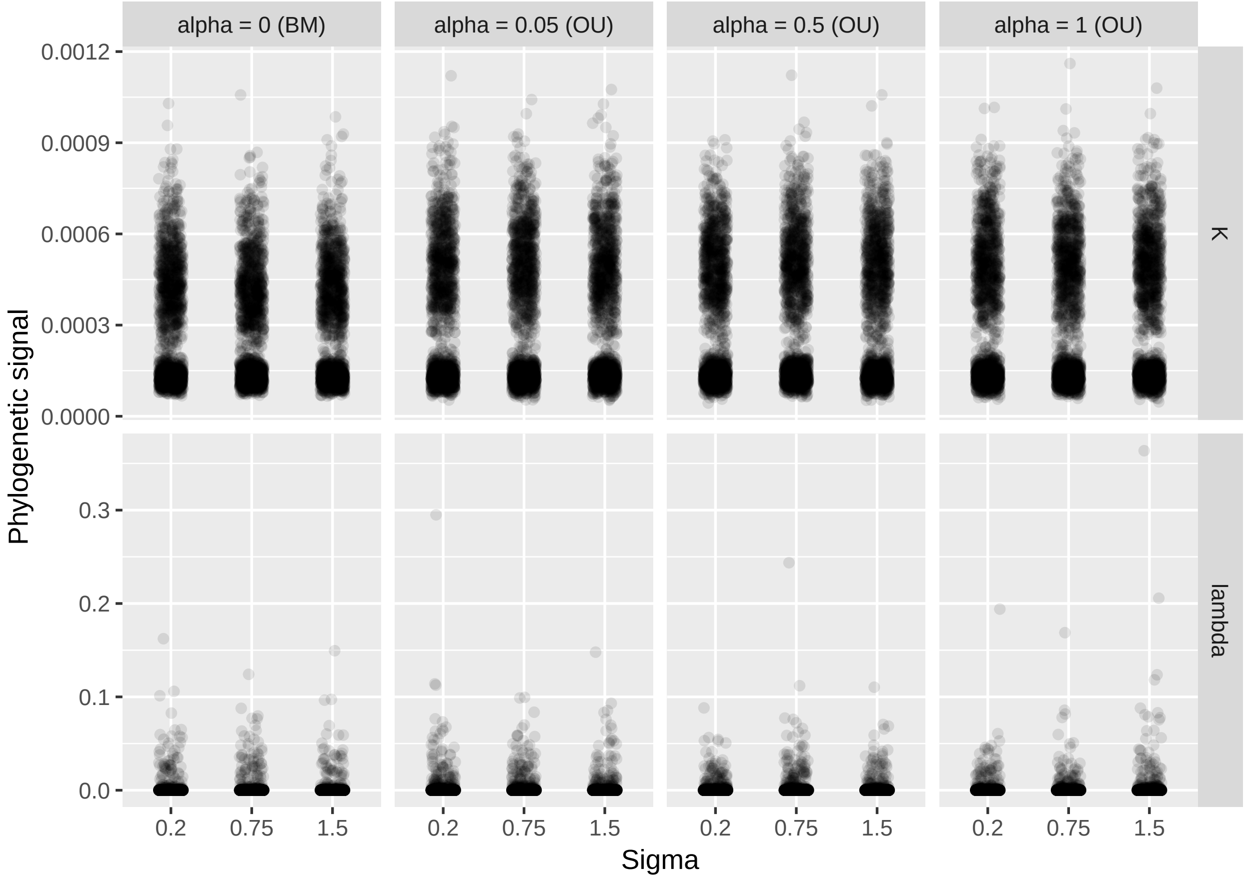


FIGURE **S7**: Estimated phylogenetic signal values based on `tree_random` were all around zero as expected.

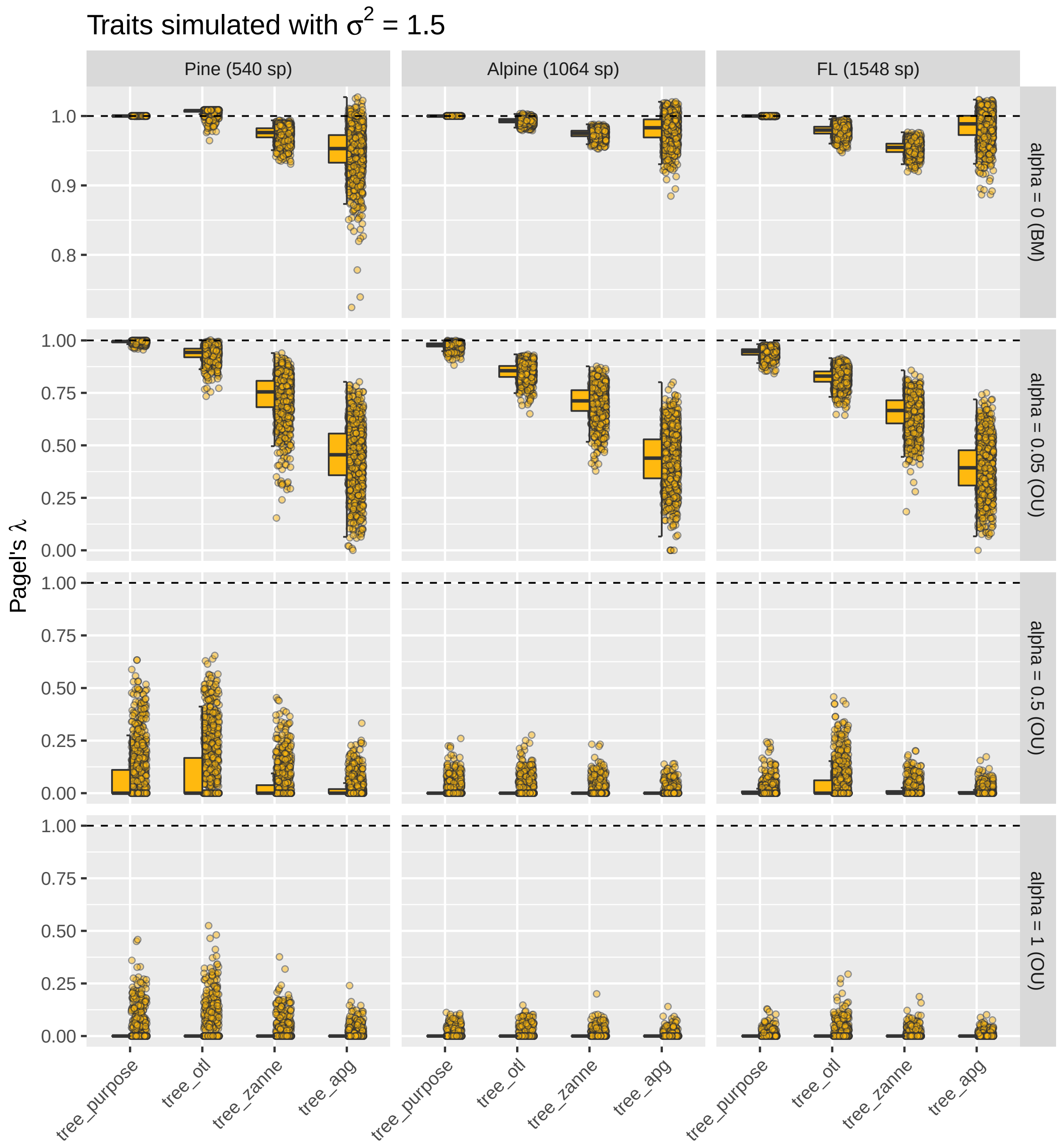


**Figure S8**: Estimated Pagel's λ when traits were simulated with divergence rate $\sigma^2$ = 1.5. The patterns here were similar as those in Fig. 4 in the main text, suggesting that sigma did not affect the estimates of Pagel's λ.

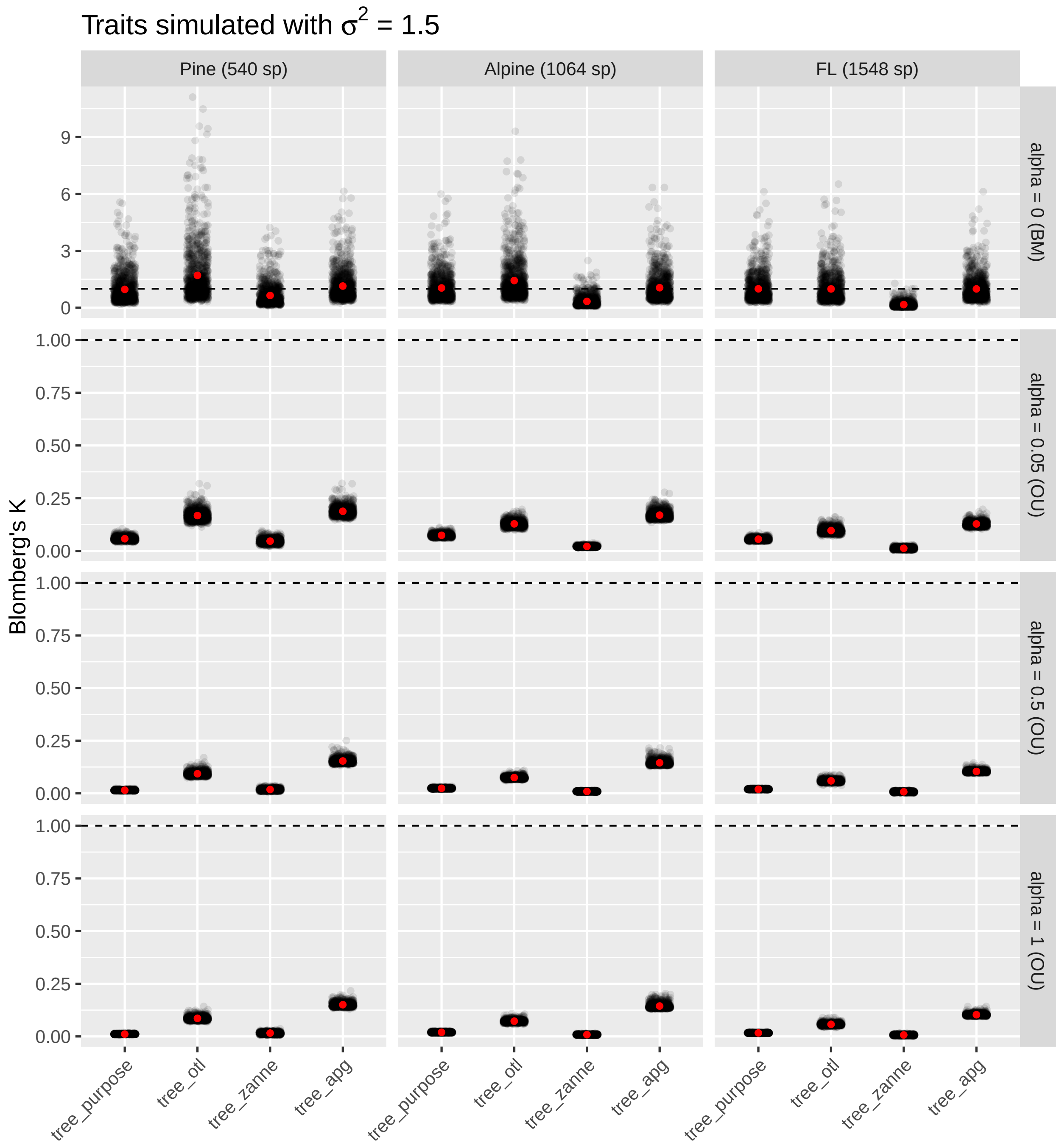


FIGURE **S9**: Estimated Blomberg's K when traits were simulated with divergence rate $\sigma^2$ = 1.5. The patterns here were similar as those in Fig. 5 in the main text, suggesting that sigma did not affect the estimates of Blomberg's K.

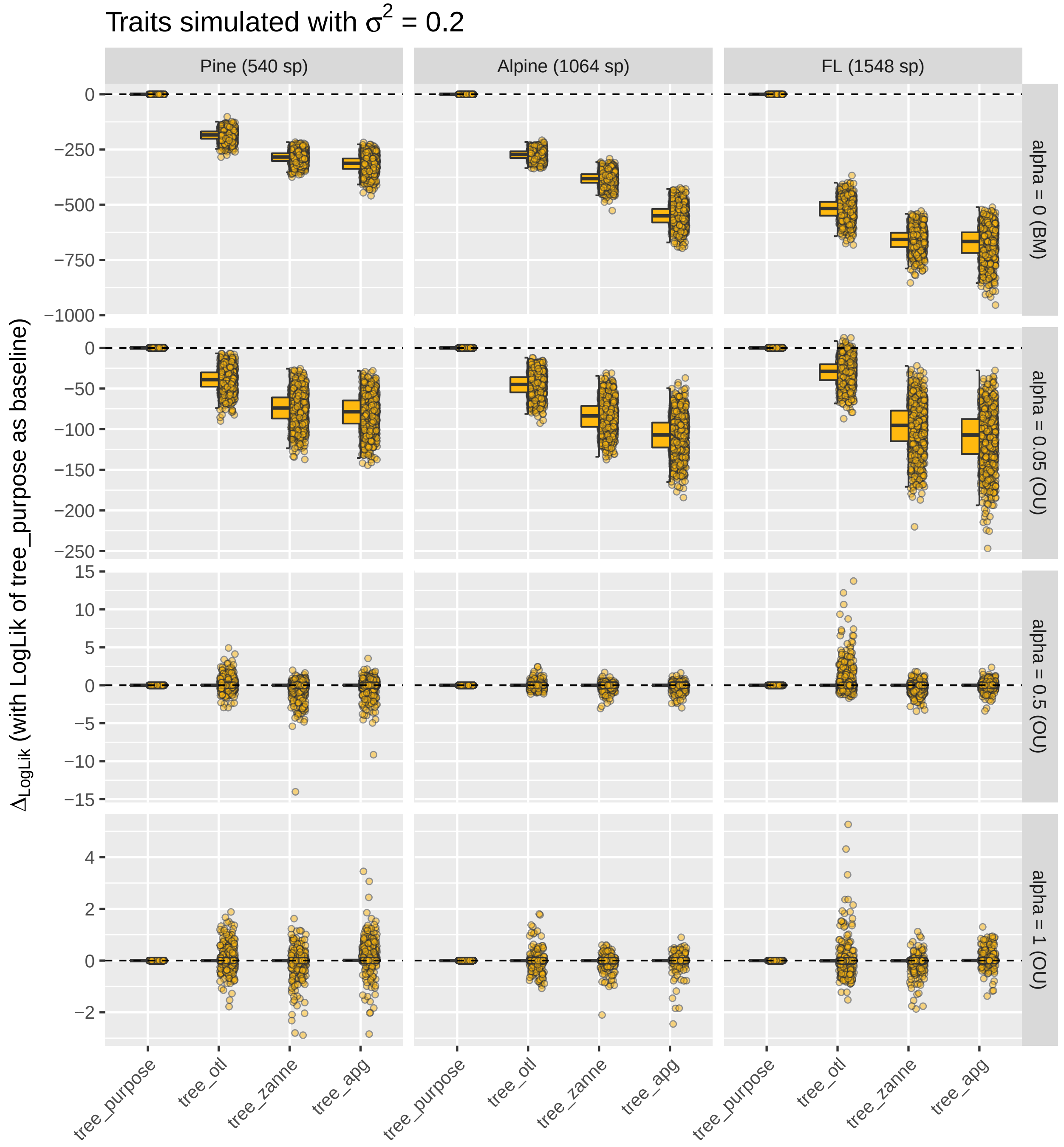


Figure S10: Differences in Log likelihood (LogLik) between `tree_purpose` and other synthesis phylogenies when estimating phylogenetic signal with Pagel's λ. LogLik values of `tree_purpose` were used as the baselines. `tree_otl` has the highest Loglik among all synthesis phylogenies when traits were simulated under BM (alpha = 0) or weak OU (alpha = 0.05). When traits were simulated under strong OU (alpha = 0.5 and 1), all phylogenies had similar LogLik values that were not different from those of null models (i.e., no phylogenetic signal). Different divergence rates ($\sigma^2$) had the pattern thus we only plotted $\sigma^2 = 0.2$ here.

Table S1: The proportion of significant estimates ($p < 0.05$) of phylogenetic signal (Pagel's λ and Blomberg's K) for traits simulated under different models of evolution with $\sigma^2 = 0.2$: BM (alpha = 0), OU (alpha = 0.05, 0.5, and 1), and random (i.e., alpha = Infinity). For traits simulated under BM, the proportion of significant results quantifies statistical power. For traits simulated under OU, the proportion of significant results quantifies statistical power in detecting less-than-BM signals (note the much higher power of K than λ when alpha = 0.5 and 1 for `tree_purpose`). For traits simulated randomly, the proportion of significant results quantifies type I error rate (i.e., false positive). Abbreviations: apg, `tree_apg`; otl, `tree_otl`; purpose, `tree_purpose`; zanne, `tree_zanne`.

| | | λ | | | | K | | | |
|---|---|---|---|---|---|---|---|---|---|
| data_set | true_alpha | apg | otl | purpose | zanne | apg | otl | purpose | zanne |
| Pine (540 sp) | 0.00 | 1 | 1 | 1 | 1 | 1 | 1 | 1 | 1 |
| | 0.05 | 0.977 | 1 | 1 | 0.991 | 0.876 | 1 | 1 | 0.999 |
| | 0.50 | 0.070 | 0.115 | 0.100 | 0.046 | 0.073 | 0.342 | 1 | 0.196 |
| | 1.00 | 0.025 | 0.029 | 0.027 | 0.016 | 0.064 | 0.152 | 0.994 | 0.105 |
| | Inf | 0.004 | 0.002 | 0.003 | 0.003 | 0.038 | 0.045 | 0.036 | 0.042 |
| Alpine (1064 sp) | 0.00 | 1 | 1 | 1 | 1 | 1 | 1 | 1 | 1 |
| | 0.05 | 0.988 | 1 | 1 | 1 | 0.486 | 1 | 1 | 1 |
| | 0.50 | 0.010 | 0.022 | 0.020 | 0.009 | 0.053 | 0.136 | 1 | 0.213 |
| | 1.00 | 0.001 | 0.010 | 0.006 | 0.005 | 0.056 | 0.073 | 1 | 0.084 |
| | Inf | 0.002 | 0.000 | 0.001 | 0.001 | 0.043 | 0.042 | 0.048 | 0.049 |
| FL (1548 sp) | 0.00 | 1 | 1 | 1 | 1 | 1 | 1 | 1 | 1 |
| | 0.05 | 0.998 | 1 | 1 | 1 | 0.965 | 0.998 | 1 | 0.585 |
| | 0.50 | 0.069 | 0.123 | 0.077 | 0.059 | 0.071 | 0.107 | 0.998 | 0.066 |
| | 1.00 | 0.016 | 0.023 | 0.017 | 0.009 | 0.051 | 0.053 | 0.631 | 0.074 |
| | Inf | 0.005 | 0.002 | 0.005 | 0.004 | 0.031 | 0.044 | 0.051 | 0.051 |
